# Event-Related Variability is Modulated by Task and Development

**DOI:** 10.1101/2021.03.07.434162

**Authors:** Shruti Naik, Parvaneh Adibpour, Jessica Dubois, Ghislaine Dehaene-Lambertz, Demian Battaglia

**Affiliations:** Cognitive Neuroimaging Unit U992, NeuroSpin Center, F-91190 Gif/Yvette, France; Université de Paris, NeuroDiderot, Inserm, F-75019 Paris, France; Institute for Systems Neuroscience U1106, Aix-Marseille Université, F-13005 Marseille, France; University of Strasbourg Institute for Advanced Studies (USIAS), F-67000 Strasbourg, France

## Abstract

In carefully designed experiments, cognitive scientists interpret the mean event-related potentials (ERP) in terms of cognitive operations. However, the huge signal variability from one trial to the next, questions the representability of such mean events. We explored here whether this variability is an unwanted noise, or an informative part of the neural response. We took advantage of the rapid changes in the visual system during human infancy and analyzed the variability of visual responses to central and lateralized faces in 2-to 6-month-old infants and adults using high-density electroencephalography (EEG). We observed that neural trajectories of individual trials always remain very far from ERP components, only moderately bending their direction with a substantial temporal jitter across trials. However, single trial trajectories displayed characteristic patterns of acceleration and deceleration when approaching ERP components, as if they were under the active influence of steering forces causing transient attraction and stabilization. These dynamic events could only partly be accounted for by induced microstate transitions or phase reset phenomena. Furthermore, these structured modulations of response variability, both between and within trials, had a rich sequential organization, which, in infants, was modulated by the task difficulty. Our approaches to characterize Event Related Variability (ERV) expand and reinterpret classic ERP analyses, making them compliant with pervasive neural variability and providing a more faithful description of neural events following stimulus presentation.

## INTRODUCTION

Since Wundt (1832-1920), the purpose of psychology has been to decompose complex cognitive functions into simpler processes, or mental operations, that could be studied in relative isolation thanks to the careful manipulation of experimental parameters (Posner and DiGirolamo, 2000). Following this ambition, thousands of studies have been published each year in which the peaks and troughs of average, stimulus-locked neural time-series (i.e. Event-Related Potentials: ERPs) have been explained as neural correlates of cognitive operations. It is indeed quite remarkable that averaging neural signals across multiple presentations of the same stimulus recovers robust and reproducible responses across participants. The ERP literature has progressively identified specific neural components whose latency and scalp-topography have been related to particular cognitive operations, from sensory processes (e.g. recognition of faces: N170)(Ghuman *et al*., 2014), to high-level processes (e.g. detecting lexicon incongruencies: N400)(Kutas and Federmeier, 2000), or monitoring our own behavioral errors (ERN: Error Related Negativity)(Dehaene, Posner and Tucker, 1994).

In this framework, the ongoing/background activity is considered as an unwanted noise discarded through the averaging process (Jasper, 1937). While measurement errors and artefacts are indeed unwanted (Verleger, Gasser and Möcks, 1982), the trial-by-trial variation of the recorded signal could also be a genuine property of the participant’s brain. Furthermore, since complete cognitive processes take place within each individual trial, and mental operations can vary from one trial to the next (e.g. stimulus visibility at threshold, confidence variation, change of strategy, etc), the signature of these operations should be detectable within individual trials –without averaging. This methodological tour-de-force is sometimes accomplished by powerful time-series pre-processing or machine learning algorithms (Jung *et al*., 2001; Vahid *et al*., 2020). However, all these methods implicitly assume that the pertinent ERP is a weak signal sunk in uncorrelated noise. Is this tenet itself as straightforward as it seems?

An increasing number of studies suggest that the background activity fluctuations are part of the cognitive process itself and can bias perceptual reports and affect stimulus detection (Hesselmann *et al*., 2008; Sadaghiani, Hesselmann and Kleinschmidt, 2009). Specifically the oscillatory components of the background activity, notably in the alpha band (8-12 Hz), are long known to be suppressed at stimulus presentation (Adrian and Matthews, 1934) and whose pre-stimulus power inversely correlate with behavioral performance (Van Dijk *et al*., 2008). Nonetheless, these oscillations are never completely suppressed and such “ongoing oscillations” display rich phase dynamics that plays an important role in top-down cognitive processes (Palva and Palva, 2007) and contributes to the detection of the ERP itself (Hanslmayr *et al*., 2007). Moreover, post-stimulus activity and ongoing fluctuations do not simply add up but nonlinearly interact (He, 2013) explaining the resulting perception (VanRullen *et al*., 2011). Finally, similarities between spontaneous and stimulus-related activity increases along development (Kenet *et al*., 2003), possibly suggesting that such spontaneous activity encodes the structure of the environment as priors (Berkes *et al*., 2011; Pezzulo, Zorzi and Corbetta, 2021). In such an alternative framework, brain activity is thought to be sampling a high-dimensional space of possible neural configurations. Such brain activity is considered to unfold along trajectories that are seemingly erratic and stochastic, and yet are loosely shaped by a latent “dynamical landscape” defined by attractor valleys and ridges connecting them (Mazor and Laurent, 2005; Gu *et al*., 2018; Chaudhuri *et al*., 2019). Spontaneous activity can thus organize in reproducible “microstates” which are visited in complex sequences, differing from mere random walks (Van de Ville, Britz and Michel, 2010). This irregular activity can still be modulated by the task demands, arousal, vigilance, etc. at the moment of stimulus presentation (Huk, Bonnen and He, 2018).

Compatible with this scenario, it was observed that inter-trial variability (which reflects the background activity fluctuations) is not constant but is characteristically reduced in the post-stimulus period with respect to baseline at rest. This “variability quenching”(VQ) after stimulus presentation is a cortex-wide phenomenon robustly observed at many spatiotemporal scales and across many different tasks (Churchland, Yu, *et al*., 2010; Arazi, Censor and Dinstein, 2017). Although different mechanisms may be responsible for it at different scales –e.g. change in excitatory/inhibitory synaptic currents at the micro-scale (Hennequin *et al*., 2018), or power increase or phase reset of ongoing oscillations at the macro-scale (Daniel *et al*., 2019; Iemi *et al*., 2019)–, the net functional effect in all cases is similar and corresponds to an increased reproducibility of neural trajectories, which, in human adults, can be further improved by active attention (Broday-Dvir *et al*., 2018) or conscious awareness (Schurger *et al*., 2015). Further evidence is nevertheless needed to understand whether this variability reduction is an epiphenomenon or plays a direct role in information processing. We argue that if variability modulations are functionally important (rather than noise), they should have a temporal structure as is the case with ERPs and this structure should emerge relatively early in life. Moreover, this phenomenon might get progressively more complex along development, reflecting the scaffolding of perceptual and cognitive processes.

Here we sought to understand the organization of response variability by reanalyzing high-density EEG data (128 channels) obtained in 5 to 24-week-old human infants (Adibpour, Dubois and Dehaene-Lambertz, 2018) as well as in adults when they were presented with human faces. We chose to study this question in human infants for three reasons: Firstly, because during the first semester of life, rapid and inhomogeneous maturation takes place, especially in the visual domain (Braddick and Atkinson, 2011), allowing age to be used as a factor to separate different neural/cognitive processes that might overlap in already mature adult brains. The peripheral visual structures reach maturity during the first semester, accompanied by a rapid myelination of optical radiations and synaptogenesis in primary visual areas. This leads to a remarkable acceleration in the latency of ERP component P1, from around 350 ms at birth to 100ms (the adult value) around 12 weeks(McCulloch, Orbach and Skarf, 1999). Interestingly, the left and right hemispheres do not mature at the same rate(Chiron *et al*., 1997), in the motor, language (Dubois *et al*., 2009) or visual system (Adibpour, Dubois and Dehaene-Lambertz, 2018) allowing for a direct comparison of the impact of maturation on similar neural pathways as a function of the hemifield of stimulus presentation. The second reason is the observation that human infants are exceptional learners (Ghislaine Dehaene-Lambertz and Spelke, 2015). If variability modulation is an intrinsic part of the building and manipulation of internal models (Berkes *et al*., 2011) the fast learning pace of infanthood might reveal more complex dynamical changes than the adults who possess relatively stable internal models. Finally, it is a common belief to disregard ongoing activity as a nuisance that compromises the robustness and reproducibility of infant ERPs. We might thus miss important information on the potential structure of the variability modulation in single-trial responses that might lead to better hypotheses and tools to gauge infant cognition.

We derived three novel measures based on multivariate pattern analysis to track the single trial dynamics and variability induced by the visual stimuli: First, we sought to quantify how individual trial trajectories approach the well-known ERP components (referred hereafter as “*ERP flybys*”). This allowed us to evaluate how the single-trial distributions of latency and distances to the ERP templates develop during the first semester of life in comparison to adults. Secondly, we examined the *“between-trial variability”* to quantify how close (or far) individual trial trajectories remained from each other as they evolved through time. This corresponds to the variability quenching phenomenon described earlier. Thirdly, we introduced a novel metric of instantaneous rate of brain state reconfiguration i.e. “*within-trial speed*” to track the moment-to-moment fluctuations along individual trials. Finally, as activity fluctuations have oscillatory components, we also studied how the dynamics of the three metrics above relate to alpha oscillatory dynamics and, specifically to alpha phase reset, since stimulus-induced alpha phase reset has been proposed as one of the mechanisms for variability quenching (Iemi *et al*., 2019).

Our results reveal a surprisingly complex temporal organization of response variability in our young participants. This organization gradually emerged through early infancy, and by the second trimester of life, reached a spatiotemporal structure remarkably similar to that of adults. Moreover, these characteristic sequences of variability modulations in infants were task-dependent. While many measures were modulated by age, we also observed in the same infants, differences in variability modulation depending on the perceptual difficulty of the task -- central faces being easier than lateralized faces. Importantly, the variability of trajectories across different trials remained daunting in both phase and amplitude of alpha oscillations even during the events of stronger variability quenching. Ongoing rest fluctuations of activity were never suppressed, as we could track by identifying microstate transitions (Michel and Koenig, 2018) and showing that their rate and probability of occurrence are barely affected by stimulus presentation. Visual stimuli neither exerted a complete reset of the system toward specific positions in the high-dimensional space of possible configurations, nor constrained the trajectories to follow specific paths with precision. On the contrary, the effects of stimulus presentation were “modulatory”, slightly bending the trial trajectories towards specific directions and accelerating or decelerating the speed of topography reconfiguration at precise post-stimulus latencies. In other words, the stimulus did not impact “where the system is” as much as it impacted “how the system flows” after stimulus presentation.

Classic ERP components were approached by single trials with very large distances and with conspicuous temporal jitter. Yet, they acted as “attractors” or “repellers” for system’s trajectories, exerting biasing forces, visible in the acceleration and deceleration patterns of within-trial reconfiguration, getting more marked and structured as development progresses. These speed modulations tended to be phase-locked to transient alpha oscillations, however the complexity of their spatiotemporal organization could not be fully accounted by phase reset events alone. We propose the term *Event-Related Variability* (ERV) to collectively describe this remarkable sequential and task-specific organization of flybys, variability quenching and boosting events, both between and within trials, which complements the classic descriptions of the modulations of average response (ERP). Such nontrivial ERV dynamics reveals an immediate richness of structured states in infants comparable to adults, confirming a potential role of variability modulations as a computing resource since the earliest ages.

## RESULTS

### 1. Event-Related Potentials (ERP) evoked by face presentation in infants and adults

Both infants (N=39, 5-24 weeks) and young adults (N=13, 21-27 years) were presented with unfamiliar faces, alternatively between the lateral hemi-fields, and for a subset of infants (N=22, 5-22 weeks), separately in the central visual field (Fig 1A, Adibpour, Dubois and Dehaene-Lambertz, 2018). Classical ERP analyses revealed two prominent ERP components: an early “P1” and a late “P400”. These components, commonly identified in infants in response to visual images and particularly faces, correspond to different cognitive stages: P1 is considered as the first cortical response in primary visual areas whereas the P400 is a higher order response related to face perception and stimulus familiarity, with sources in the fusiform region (De Haan, Johnson and Halit, 2003). These components, visible in the grand average topography in Fig 1B for infants, also had clear equivalent topographies in adults (Fig S1A). For adults, latencies were faster, voltage topographies qualitatively similar and overall ERP signal amplitude weaker as compared to infants. For lateral faces, the P1 response corresponded to the first positivity on the contra-lateral posterior electrodes around 250-300 ms following face onset in infants (∼100 ms in adults). The P400 response was a large bilateral positivity on occipitotemporal clusters following the P1 response around 500-600 ms (∼400 ms in adults). For central faces, latencies were faster relative to lateralized faces: around 150 ms for the P1 visible on medial posterior electrodes and around 450-550 ms for the P400 on occipitotemporal channels (Fig 1B bottom row). The overall signal amplitude was larger for central than for the lateral faces. These results are in agreement with previous literature on ERP dynamics following face presentation in adults and infants (De Haan, Johnson and Halit, 2003; Conte *et al*., 2020)(Conte et al., 2020; De Haan et al., 2003). Infant N290 topography was not prominent and adult N170 topography did not find clear equivalence in infants, hence we avoided these “intermediate” components.

**Fig 1.**
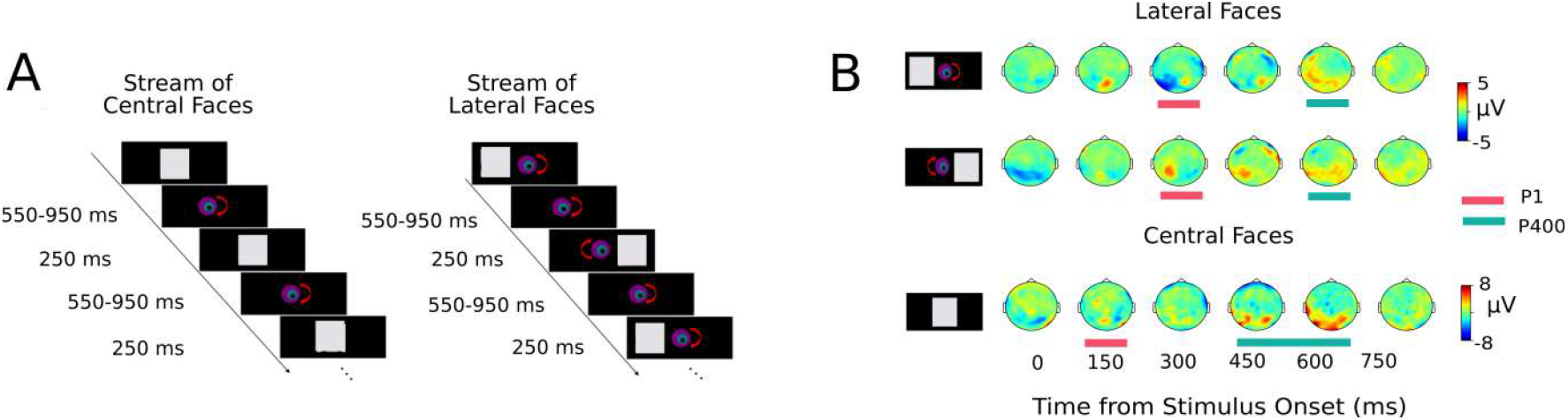
Task Paradigm and Infant visual ERPs. **A)** Infants (and adults) were presented with unfamiliar faces consecutively in the left and right hemi-field. A subset of infants was also presented with faces in center. **B)** Grand Average voltage topographies for the three conditions for infants. Early (P1) and late (P400) ERP components (marked with red and green horizontal bars respectively) are visible for each condition. Human Face stimuli are replaced by gray boxes (here and elsewhere) in compliance with the Biorxiv privacy policy.

### 2. Ongoing Variability Dominates ERP

While characteristic ERPs existed, single-trial responses were noisy and hardly resembled grand averaged responses (Fig 2A; Fig S1B), typically remaining one- or two-order of magnitude larger than ERP amplitudes in both adults and infants, with extremely variable topographies and no clear peaks or valleys corresponding to the ERP.

**Fig 2.**
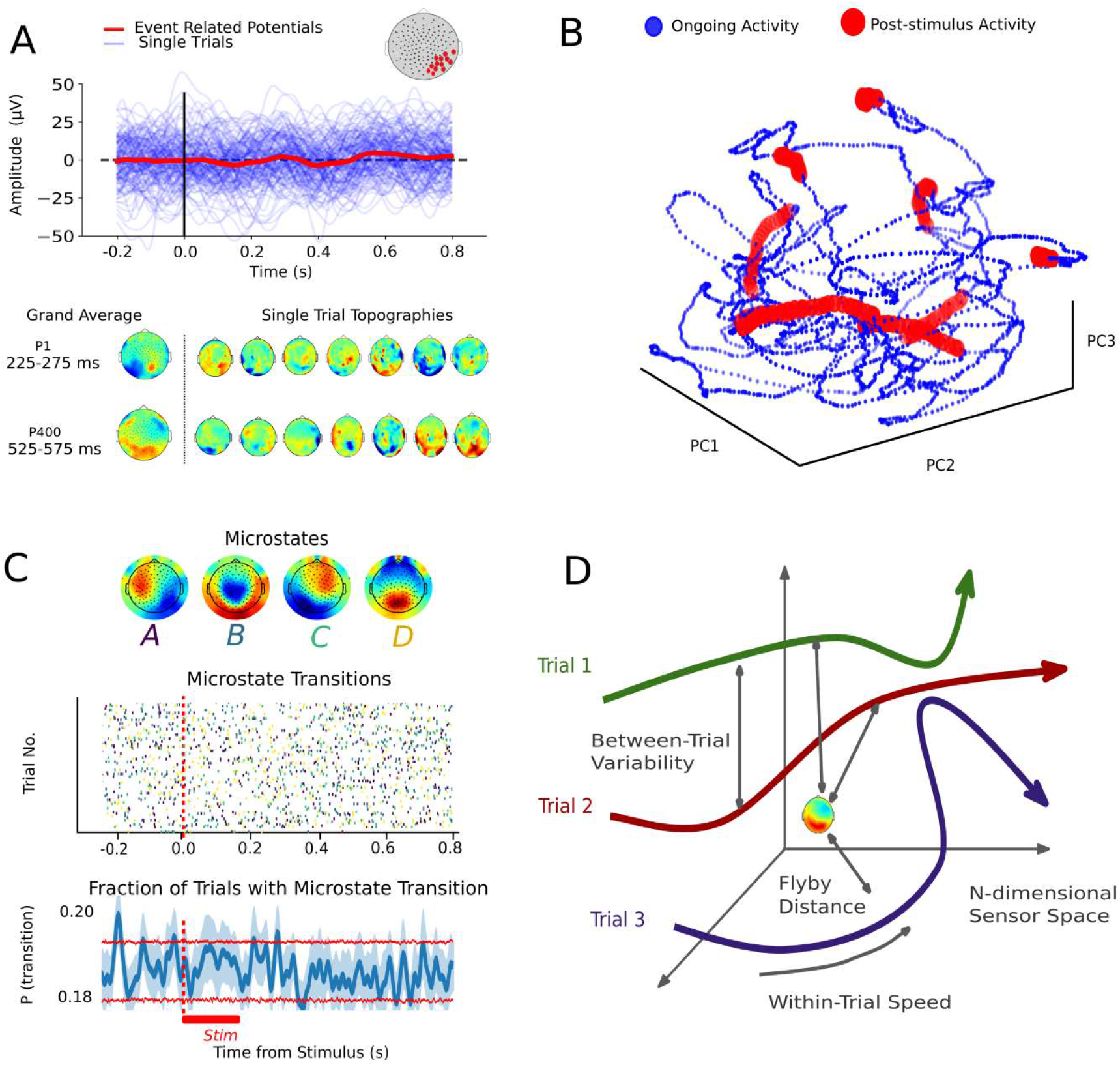
Ongoing Variability dominates Event Related Potentials. **A)** Example voltage time-courses averaged across right occipital electrodes for a representative infant (age =12.1 weeks) for faces presented in left hemi-field. Amplitude and latency of single-trial responses (blue lines) are highly variable in comparison with average ERP response (red bold line). Single-trial voltage topographies in P1 and P400 response range are notably different from the grand average ERP topography. **B)** Trajectory of continuous time-segment (∼ 12 s) reduced to 3-dimensional PC space from 128-channel EEG sensor space for the same infant. Each point corresponds to single instantaneous voltage topography. Time-points falling in the 450-500 ms post-stimulus time range are marked in red. **C)** (Top) Topographies of infant microstates. (Middle) Raster plots of single-trial microstate transition trains. E ch dot marks a change in microstate at that time point. The color of dots indicates which microstate was transitioned into at that time point (corresponding to the colors of letters indicated below the microstate topographies). 100 randomly chosen trials for a representative infant are plotted for visibility. No clear pattern of transitions is visible. (Bottom) Gaussian smoothed microstate transition rates averaged across all trials and infants (see Methods). Red horizontal lines represent [5-95] % confidence interval. The absence of peak in this curve shows the microstate dynamics remain unperturbed. **D)** Schematic summary of methods used to gauge single-trial variability: Flyby to ERP templates (how far a trial transits from reference ERP-like configurations); Between-trial variability (how far are single trial trajectories between them); and, Within-trial speed (how fast EEG topographies evolve along each trial).

To illustrate the relationship between the ongoing and evoked activity, we show a 12 s-long-segment of continuous EEG data for a representative infant (age=15.4 weeks) in Fig 2B. At every time point, brain topography is represented as a point in a three-dimensional projection after dimensionality reduction (performed by applying Principal Component Analysis (PCA) to activity amplitudes, see *methods*). If stimulus presentation always evoked a similar single-trial trajectory, the EEG trajectory time points recorded immediately following the presentation of a face stimulus should cluster together within this low-dimensional projection. On the contrary, post-stimulus trajectory snippets (highlighted in red in Fig. 2B) were distributed nearly uniformly throughout the sampled space. This dispersion suggests that post-stimulus temporal fluctuations of neural trajectory were predominantly determined by the ongoing activity. Individual stimulus presentation events seemed not to lead to a radical reset of the activation topographies and did not constrain them to strongly resemble the grand-average topographies.

Ongoing spontaneous activity has been shown to have a rich spatiotemporal organization, which has been previously characterized in terms of discrete ‘Microstate transitions’ (da Cruz *et al*., 2020). Microstates are patterns of scalp topographies extracted through unsupervised clustering procedures and are transiently stable for ∼60–120 ms in adults. They are shown to stochastically alternate between themselves during spontaneous resting activity, with a scale-free distribution of their dwell-times (Van de Ville, Britz and Michel, 2010). Here we extracted microstate sequences from EEG recordings as a possible, simple way to apprehend the organization of global ongoing fluctuations. We describe them as transitions between a set of unsupervised reference topographies, which can be thought as an average orientation frame (without necessarily committing on their exact number or spatial pattern). We checked whether presentation of the stimulus led to a perturbation of ongoing dynamics, in terms of enhanced probabilities to observe certain microstates or stimulus-triggered microstate transitions (see *Methods*). Fig 2C (top) shows the 4 microstates extracted by the standard k-means clustering of the continuous EEG time segments across all infants during the lateralized face paradigm. The topographies were reminiscent of the ones commonly observed for adults in microstate studies. In Fig 2C (middle panel), we visualize the transitions from one microstate to another as a raster plot, to understand whether the transition probabilities were modified by the stimulus presentation. If microstate transitions became more frequent at a certain fixed post-stimulus latency, one would observe the formation of vertical stripes in this raster (depicting time-aligned transition for each trial) possibly of relatively uniform color (indicating a specific microstate being associated to post-stimulus activity). On the contrary, the raster plot appears unstructured and “asynchronous”, with a salt-and-pepper arrangement of colored dots, denoting that stimulus presentation does not induce the selection of a specific microstate. This is quantitatively confirmed by Figure S2 in which we show the near-absence of modulations in the probability of visiting specific microstates as a function of stimulus onset time (with mild variation across age-groups). We then quantified whether stimulus presentation modifies the rate of generic microstates transitions, irrespectively of microstate identity. The time-course of this average microstate transition rate, shown in Fig. 2C (bottom panel), reveals once again a complete absence of significant modulations of microstate transition dynamics. Spontaneous ongoing fluctuations continued to occur in an unaltered fashion, as if virtually unaffected by stimulus presentation.

These analyses suggest that the ongoing dynamics is not radically reconfigured or perturbed by stimulus presentation in its global aspects. On the contrary, stimulus-induced effects appear to be riding on top of ongoing fluctuations. System’s trajectories can be virtually “everywhere” in the high-dimensional space of possible topographies and stimulus-related effects must correspond to small local amplitude or phase modulations of current trajectories rather than a radical channeling of the trajectories along specific paths.

With the hypothesis that this tremendous response variability and moment-to-moment fluctuations is informative about underlying neural and cognitive processes, we considered three indicators in order to detect weak stimulus-related modulations on top of highly variable signals (see cartoon representations in Fig. 2D) as previously described: *ERP component flybys, between trial variability* and *within-trial variability (or trial Speed)*. To quantify (dis-)similarity between topography patterns, occurring in different trials or at different times along a trial, we used a multivariate metric of correlation distance, i.e. 1 – Pearson Correlation Coefficient (see *Methods*). Such a measure is sensitive to variations of the relative strengths –or activity patterns– rather than to the absolute voltage amplitudes observed at specific EEG channels. For this reason, we believe it is suitable to identify weak local reconfiguration events that ride on top of simultaneously ongoing global fluctuations. Finally, we verified the relation between the dynamics of our metrics of variability with the phase and amplitude of the signals filtered in a narrow alpha band (Methods), to study their eventual relation with phase reset events. Results from all the three approaches are summarized and put in relation with alpha oscillation dynamics in the following sections.

### 3. Single-trial flybys to classic ERP components are modulated by age and hemisphere

Despite erratic trajectories, grand average ERP topographies are reproducible across studies suggesting they capture stimulus-relevant information. Hence, we analyzed how individual trials approach (or flyby) these “landmark events”. We defined the grand averaged P1 and P400 topographies (separately for infants and adults) as “ERP-templates” (Fig S3) and examined the distributions of the latencies and distances of the single-trial flybys to these templates (Fig 3 and S4, see *Methods*).

**Fig 3.**
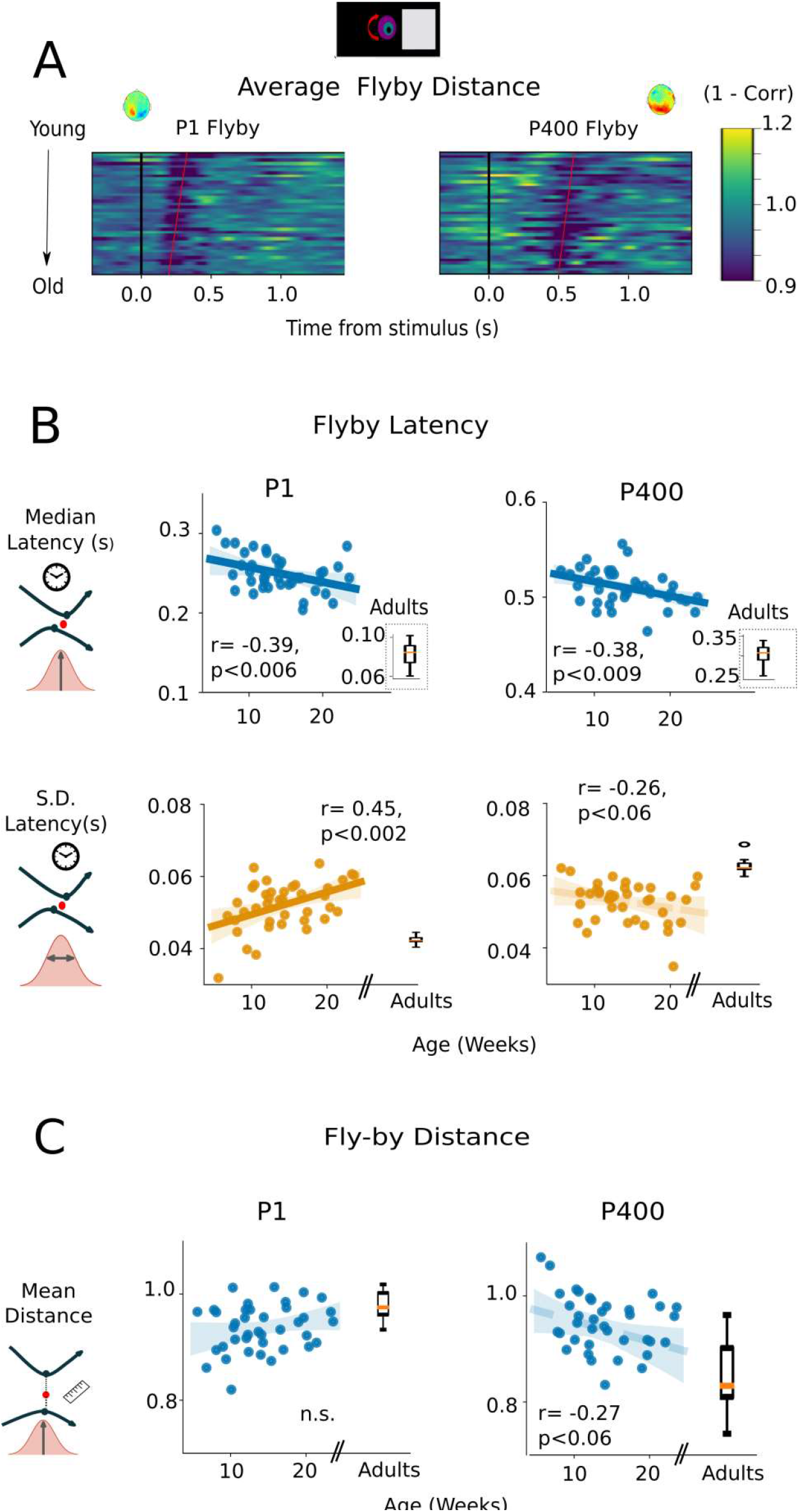
Maturation of single-trial Flyby statistics for faces presented on the right hemi-field. **A)** Average flyby distances to P1 (left) and P400 (right) ERP templates (shown on top) for each infant. Each row represents a single infant, sorted in ascending order according to their age (from youngest = 5.6 weeks to oldest=23.1 weeks). Red vertical lines emphasize the reduction in average flyby distance from ∼150-350 ms for P1 and ∼400-600ms for P400 templates. The slopes of red lines suggest that latency of closest distance reduces with age. **B)** Median flyby latency significantly decreased with age for both P1 (top left) and P400-flyby (top right panel). At the same time, S.D. of single-trial flyby latencies significantly increased with age for P1 (bottom left panel) and showed a negative trend with age for P400 template (bottom right panel). Inset box-plots represent the same statistics for adults. **C)** Average flyby distances to P1 template showed non-significant increase with age (top left panel), while the same for P400 template decreased with age (top right panel). Boxplots indicate that once again adults followed the same trends. Shaded areas indicate 95% confidence interval for the slope of the least square fitting line, all r-values corrected for multiple comparisons with 1-tailed permutation t-test).

For all task-conditions, trials remained quite far from the ERP-templates most of the time (flyby distance ∼0.8-1; Pearson correlation: 0-0.2, Fig3A, FigS4A, B), in line with the finding that system’s trajectories fill the entire configuration space. However, individual trial trajectories slightly and significantly reduced their distance to the ERP-templates around specific latencies (Distance drops in Fig. 3A and S4A respectively for right and left faces, emphasized by red vertical lines), in line with the intuition that stimulus induces small trajectory inflections independently from where exactly the system is transiting. For lateral faces, trials approached the P1-template around [150, 350] ms; and the P400-template about [400,600] ms post-stimulus onset. For the central faces, closest approach to ERP templates occurred, within the broad ranges between −150 and 150 ms for the P1 template, and for the P400 template between 350 and 550 ms (Fig S4 B).

To quantify age-dependency of flyby latencies and latency jitters around the ERP components, we estimated the distribution of closest single-trial flybys for each infant and adult and studied whether their medians and standard deviations were correlated with age. In agreement with P1 latency measures across ages in previous studies, we found that flybys to ERP components tended to occur at earlier latencies with increasing age. However, flybys did not necessarily become more accurate, i.e. with a more precise timing at each trial. On the contrary, for some of the ERP components, the jitter in the timing at which the closest flybys occurs (dissimilarity from reference ERP component topography) increased with age, suggesting that trajectory variability is not always a “bug” to correct for, but a feature growing through development.

More specifically, for *flyby latencies*: The median of the flyby latencies to both the P1-and P400-templates significantly decreased with infant’s age for the faces presented on the right hemifield (P1: r= −0.39, p<0.006 from ∼280 ms at 5 weeks to ∼200 ms at 24 weeks; P400: r= −0.38, p<0.009 from ∼600 ms to ∼500 ms, Fig. 3A). This decreasing trend was confirmed in adults whose median flyby latencies were much faster than in infants (inset box-plots in Fig.3B: P1 median latencies across adults: 96 ± 13 ms and P400 median latencies: 312 ± 20 ms for right faces). In each infant, the dispersion around the median latency value was ∼40-60ms. Age also affected this dispersion but differently for the P1 (Fig 3B left bottom panel, r = 0.45, p<0.002) and P400 component (Fig 3B, right bottom panel, r = −0.26, p<0.055). There was an increase with age in the spread of the flyby latency distribution, suggesting that the timing of approach to the P1-template became less accurate, or more flexible, through early development. It contrasted with the inverse and moderate trend for the P400-flybys. Interestingly, adults also showed a ∼40-60 ms jitter across trials, but similar for the two components (Figure 3 B box plots: P1-latency jitters: 54 ± 4ms and P400 latency jitters: 57 ± 3 ms for right faces).

For left and central faces, we observed no effect of age on these measures in infants (Fig S3 C, D top panels). The effect of age for the right faces (processed by the left-hemisphere) was related to an initial delay in flyby latencies in the youngest infants. Thus, the catch-up relatively to the more mature right hemisphere during this period is congruent with several results showing a slower maturation of the left hemisphere compared to right (Chiron et al., 1997). Similar catch-up of the maturation of the left dorsal linguistic pathway relative to the right has also been described during the first semester post-term (Leroy et al, 2011)

Similar age-effect analyses can be performed on the *flyby distance* to the template for each trial (flyby distance amplitude distributions: Fig 3C, Supp. Fig S4 E, F). No age-effect was significant for P1 flybys. For P400 flybys, the mean distances reduced with age in the case of left, right and central faces, suggesting that on an average, trials for older infants passed closer to P400-template than those for young infants. (r = −0.37, p<0.02, r=-0.52, p<0.006, r = −0.26, p <0.06 for left, right and central faces respectively). For P400-flybys, these trends continued well into the adulthood, i.e., trials passed on average much closer to the template (Fig 3C, right box insets and Fig S4 E, F box-plots and table 1). Finally, the variability of the flyby distances either remained unchanged, for the P1-template; or even grew with age, for the P400-template (age correlation for infants, left faces: r = 0.42, p<0.003; right faces: r = 0.34, p<0.02 and central faces: r = 0.53, p<0.007, with the similar trend continuing for adults).

**Table 1.** Comparison of P1 and P400 Flyby Mean Amplitude Across Age-Groups. Table 1 Average closest flyby distances across age-groups (5-12 week old: first trimester infants, 16-24 weeks old trimester infants and adults) were compared using separate Kruskal-Wallis tests for different ERP templates (P1, P400 responses) and for faces presented in the left and right hemi-field. The main effects are reported before post-hoc Mann-Whitney U-test for pair-wise comparisons. P values are corrected for multiple comparisons using Bonferroni correction.

| Table 1 | Comparison of P1 and P400 Flyby Mean Amplitude Across Age-Groups |  |
| --- | --- | --- |
|  | Left Faces | Right Faces |
| P1 | H = 11.48, p=0.003<br>5-12wo vs 16-24 wo: n.s.<br>5-12 wo vs adults: p= 0.009<br>16-24 wo vs adults: p= 0.015 | H = 9.76, p=0.007<br>5-12 wo vs 16-24 wo: n.s.<br>5-12 wo vs adults: p=0.014<br>16-24 wo vs adults: 0.072 n.s. |
| P400 | H = 13.29 p = 0.0013<br>5-12wo vs 16-24 wo: p = 0.18<br>n.s.<br>5-12 wo vs adults: p = 0.001<br>16-24 wo vs adults: p = 0.25 n.s. | H=13.47,p=0.001<br>5-12 wo vs 16-24 wo: p= 0.4n.s.<br>5-12 wo vs adults: p= 0.003<br>16-24 wo vs adults: p =0.028 |

To summarize, single-trial event-related dynamics significantly changed with age. ERP flybys became in general more fluid (faster and relatively more variable in timing or distance of approach). The observed developmental changes to ERP flybys were specific for the considered ERP component (P1 or P400) and, in the case of infants, for the hemisphere probed by the lateralized stimulus (indicating thus a possible influence of selective connectivity maturation). In all cases and at all ages, single trial trajectories remained rather distant from ERP templates even at the moments of closest fly-by (with correlation distances larger than ∼0.8, not so far away the unit value which would correspond to complete lack of correlation), compatibly with the large variability observed earlier (c.f. Fig. 2).

### 4. Between-trial variability quenching (VQ) after stimulus presentation

Irrespective of their approach to the templates, trials can remain far or close to each other at any point. Hence, we investigated between-trial variability. Again we found that trajectories remained highly dissimilar, as denoted by an average correlation distance of 0.95±0.12 between the time-aligned trajectories of different trials. Although large in absolute terms, the between trials distance relatively reduced at specific peri-stimulus times. We observed a significant post-stimulus decrease in the between trial variability for all task-conditions and for both infants and adults (Blue plots in Fig 4A-B, sup Fig S5). For infants, between-trial variability significantly remained ∼1-2.5 standard deviations lower than the average baseline variability ∼200-700 ms post-stimulus (p < 0.001 for left, right faces and p<0.003 for the central faces), while in adults, significant Variability Quenching (VQ) occurred ∼150-500 ms (p<0.005), similar to the duration previously reported for variability quenching in adults (Schurger *et al*., 2015).

**Fig 4.**
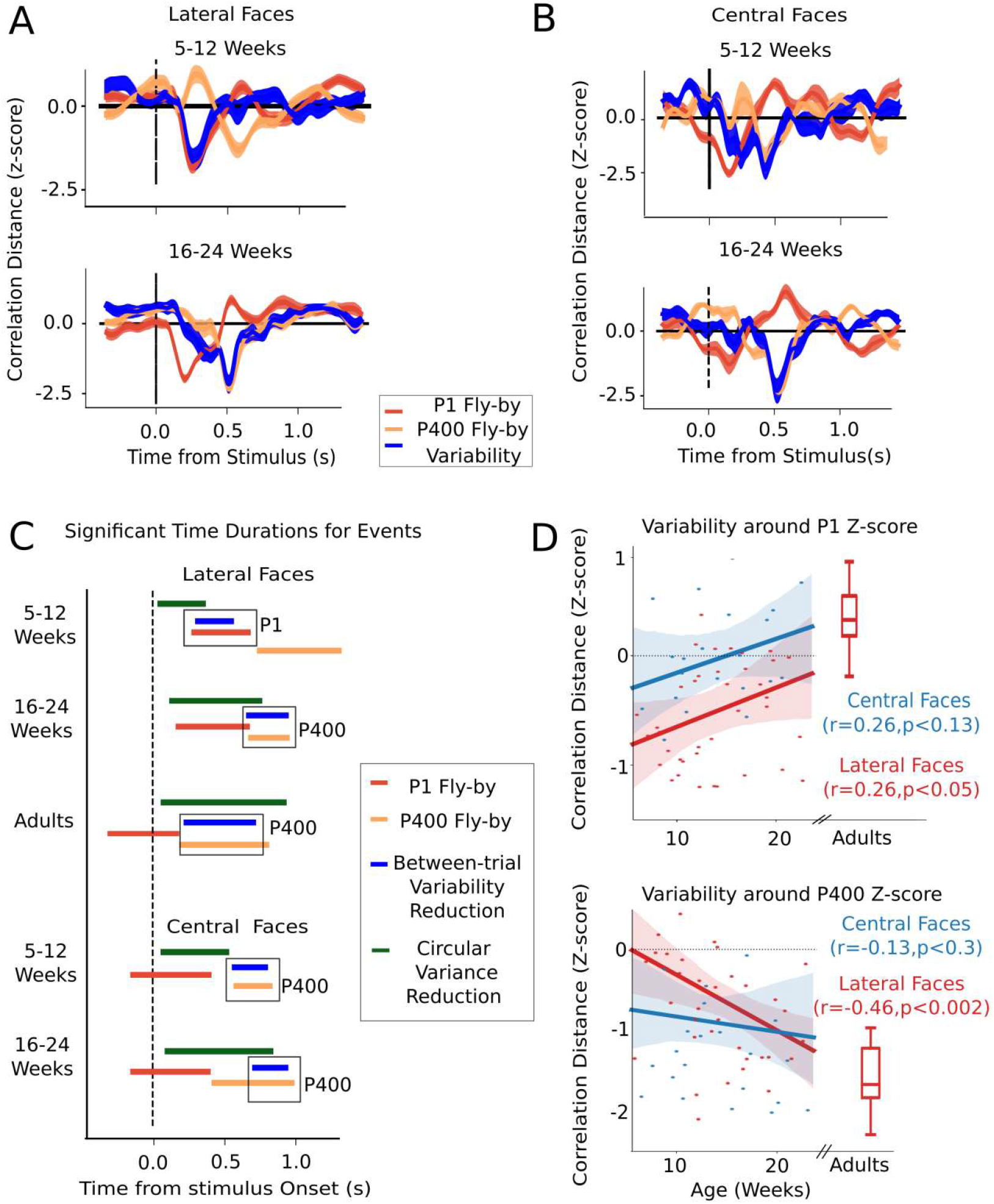
Maturation of Between-trial variability and its relation to Flyby distances. **A)** Group average between-trial correlation distance (blue curves, Z-scored) for 5-12 weeks (N= 14, top panel) and 16-24 weeks (N= 13, bottom panel) old infants plotted together with grand-average flyby distances (Z-scored) to P1 (red line) and P400 templates (orange line), for the lateral faces. Significant reduction in between-trial variability coincides with the closest-flyby to P1-template for 5-12wo infants (top panel), and wit the P400-template for 16-24wo infants (bottom panel). **B)** For central faces, significant between-trial variability (blue line) reduction coincides with P400-flyby (orange line) for both groups. Shaded areas indicate standard error to the mean. **C)** Synoptic view, across conditions and age groups, of the time ranges when between-trial variability (blue), between-trial circular variance of phases of alpha oscillations (green) and flyby distance (red /orange) are significantly reduced (Horizontal bars indicate timepoints of significant reductions from mean; p<0.05, corrected using cluster-based permutation t-test). Boxes highlight correspondence between clusters of between-trial variability and P1/P400 flyby distance reductions in different conditions.16-24 wo infants qualitatively look like adults. 5-12 wo infants when presented with central faces also quench variability at P400. In all conditions, Circular Variance (CV) reduction precedes variability quenching (VQ) events. D) Between-trial variability during P1-flyby increases with age (top) while the same during P400-flyby decreases with age for lateral but not central faces. (All r-values corrected for multiple comparison with one-tailed permutation test). Shaded area indicates 95 % confidence interval for the slope estimation of the least square fitting. Boxplots show between-trial variability distributions for adults.

#### Between-trial VQ is not automatically induced by ERP component flybys

The latency of the largest post-stimulus VQ (lowest variability) significantly differed across age-groups and task conditions. Strikingly, for lateralized faces, the latency of the significant VQ coincided with the latency of the closest P1-flyby in the youngest infants (First trimester: 5-12 week-olds, N=14) whereas in the older infants (Second Trimester: 16-24 weeks, N=13), the moments of VQ co-occurred with the P400-flyby (Fig 4A). In other words, in younger infants, the bundle of single trial trajectories remained on an average more compact when flying by the P1-template (significant VQ times: 204-352 ms, window of closest P1-flyby: 175-400 ms). By contrast, in older infants, trials remained the closest to each other when passing near the P400-template (significant VQ: 432 – 616 ms, closest P400-flyby: 432-620 ms).

Importantly, the absence of between-trial VQ did not imply absence of a flyby. Indeed, in first-trimester infants, trials still had a marked P400-flyby even when there was no between-trial VQ at the corresponding latency. Analogously, there was still a P1-flyby for second-trimester infants despite the lack of a P1 VQ. These effects were consistent for both left and right face presentation (Fig S5A-B). Thus, flying by an ERP component appears to be a necessary but not a sufficient condition for between-trial VQ. In adults too, a single window of VQ coincided with the P400-flyby, similar to the second-trimester infants’ pattern (Fig 4C, Fig S5 C, and D). However, the VQ was much larger in adults than in infants, trials remaining significantly close to each other during the entire duration of the P400-flyby (significant VQ: 140 – 460 ms, P400-Flyby: 120-528 ms).

#### Between-trials variability quenching depends on both stimulus configuration and age

The temporal shift of the between-trial VQ, from P1 to P400 gives the first proof of a change in the ongoing dynamics occurring over the first semester of life. However, such a shift may also be due to the structural changes in the peripheral visual pathway and visual cortex V1 which are known to reach a milestone around 12 weeks post-term (McCulloch, Orbach and Skarf, 1999; Braddick and Atkinson, 2011; Adibpour, Dubois and Dehaene-Lambertz, 2018). To investigate the possible origin of such a shift, we repeated the same analysis in the subset of infants who were also presented with central faces (Fig 4B). We found that the VQ at P1 but not P400-flyby significantly differed in the same infant for the two different face stimuli configurations (Wilcoxon signed rank (z)=58, p<0.025 for P1, z=123,p>0.92 for P400). Interestingly, when presenting central face stimuli, the 5-12 weeks infants now showed a significant VQ at the P400-flybys, similarly to the response to lateral faces observed in 16-24 weeks infants. The absence of VQ at P400 for lateral faces in 5-12 weeks infants thus, does not reflect uniquely a poor maturation of connection pathways. Fig 4C summarizes the time-ranges of between-trials variability quenching and P1/P400-flybys across all experimental conditions and age groups (p<0.05, all analyses were corrected for multiple comparison and temporal independence using one-sided cluster-based permutation t-test).

Age affected not only the latency but also the strength of VQ events, in similar directions for both stimulus configurations. Quenching strength decreased with age within the P1-range and increased within the P-400 range (Figure 4D), with both these trends confirming the observed inter-group differences. Specifically, we found a linear increase of the strength of between-trials variability quenching with age at the P400-flyby latency for lateral faces (Fig 4D bottom panel, r = −0.43, p <0.003 for left faces; r = −0.30, p <0.03 for right faces; but not for central faces, r = −0.13, p <0.3). Table 2 and the box plots in Figure 4D show that adult values further continue the trends observed in infancy.

**Table 2.**
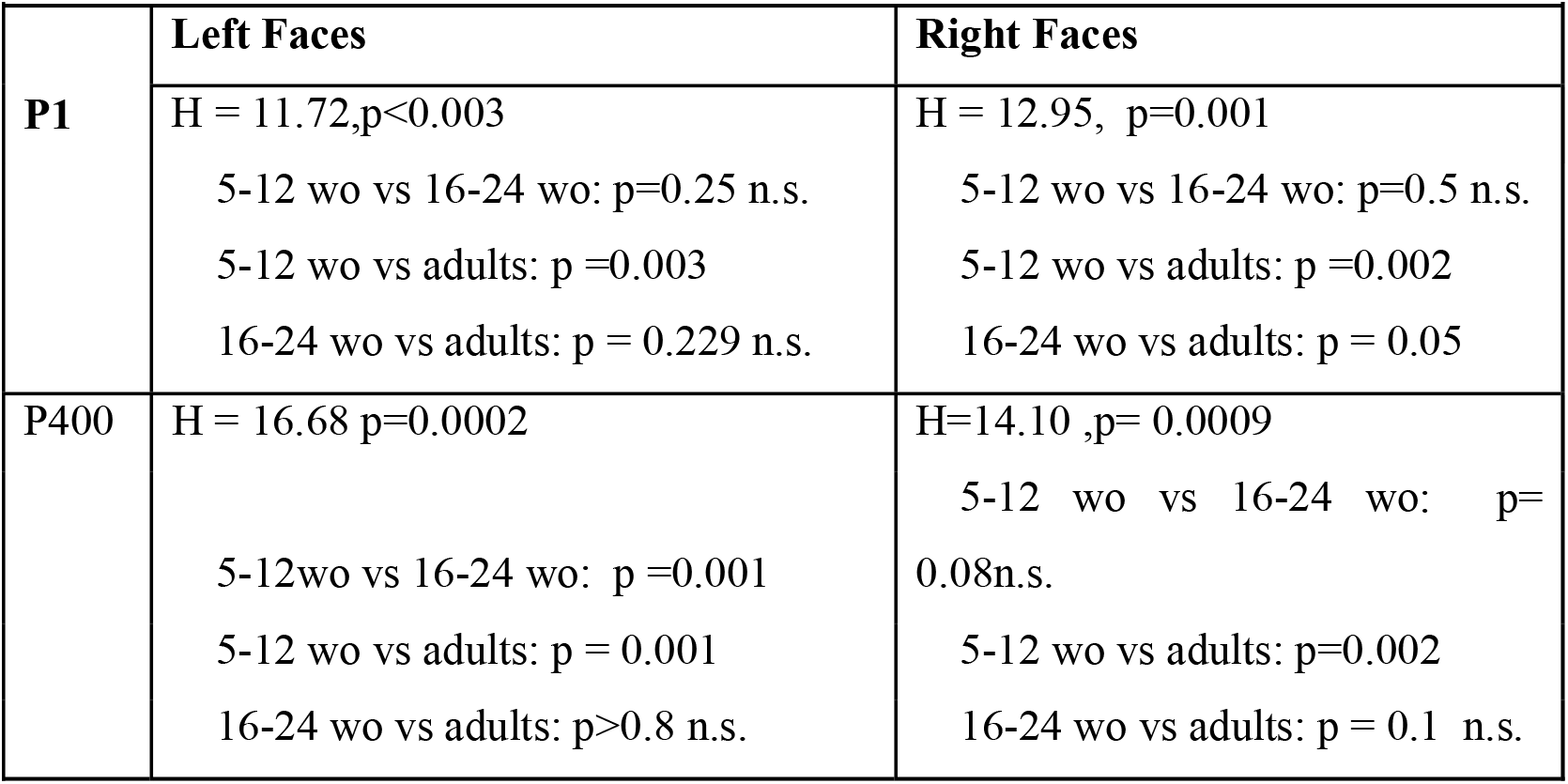
Age Difference in Between-trial Variability around Flybys (*z-scores*) Table 2 Between-trial variability (z-scores) were compared the three age-groups (5-12 week old: first trimester infants, 16-24 weeks old trimester infants and adults) in their respective moments of P1 and P400 closest flybys using separate Kruskal-Wallis tests for different ERP templates (P1, P400) and for faces presented in the left and right hemi-field. The main effects are reported before post-hoc Mann-Whitney U-test for pair-wise comparisons. P-values are corrected for multiple comparisons using Bonferroni correction.

**Table 3:** Age-differences in Within-Trial Variability Around Flybys. Table 3 Flyby triggered within-trial speed profile time-series (z-scores) across the three age-groups were compared using cluster-based permutation F-tests separately for each ERP template and for each hemi-field to find the significant time clusters (p<0.05). During post-hoc analyses, average within-trial speed in these significant time-windows was compared using Kruskal-Wallis test. Pair-wise comparisons were tested using Mann-Whitney U-test. P-values are corrected for Bonferroni correction.

| **Table 3: Age-differences in Within-Trial Variability Around Flybys** |  |  |
| --- | --- | --- |
|  | **Left Faces** | **Right Faces** |
| **P1** | S.C.: None | S.C.: [8 80] ms, H=6.07, p=0.048<br>5-12 wo vs 16-24 wo: p = 0.08<br>5-12 wo vs adults: p =0.17<br>16-24 wo vs adults: p = n.s. |
| **P400** | S.C.: [-384 -308] ms: H = 14.87, p= 0.0006<br>5-12wo vs 16-24 wo: p = n.s<br>5-12 wo vs adults: p= 0.008<br>16-24 wo vs adults: p= 0.0008 | S.C.[-400 -268]ms, H=14.42, p=0.0007,<br>5-12wo vs 16-24 wo: p = n.s<br>5-12 wo vs adults: p= 0.015<br>16-24 wo vs adults: p= 0.0005 |
|  | S.C = Significant Clusters (p<0.05, cluster based permutation tests) | S.C.[-72 -12]ms, H= 16.266, p=0.0003<br>5-12wo vs 16-24 wo: p = n.s.<br>5-12 wo vs adults: p= 0.0003<br>16-24 wo vs adults: p= 0.03 |

#### Between-trials variability quenching is not equivalent to alpha phase reset dynamics

EEG responses have oscillatory components, with a spectral resonance in the alpha band (9-12 Hz) which was relatively prominent in adult subjects, but way less marked in infants (Fig. S6A). To investigate the relation between VQ and reconfiguration of alpha oscillatory dynamics, we narrow-band filtered the EEG signals in a band of interest and extracted phase and amplitude of the oscillations through Hilbert transform (see Methods). We then quantified the time-courses of trial-averaged alpha power modulation (Fig. S6B, top) as well as the circular variance (CV) of alpha phases across time-aligned trials (Fig. S6B, middle). Average alpha power was not significantly modulated in the peri-stimulus duration in infants, it was only slightly reduced in adults (∼0.8 S.D. below the baseline) and for infants, this reduction was never significant at the level of global topography averages. The effect of stimulus presentation was slightly more pronounced on the phase of alpha oscillations. The measured CV of alpha phase across stimulus-aligned trials denoted a poor phase-alignment between-trial, with an average value of ∼0.80 (± 0.06) in infants and ∼0.93 (± 0.02) in adults, close to the unit value that would correspond to a complete asynchrony of phases across trials. Face presentation did not induce a complete reset of ongoing oscillations, once again in line with the large variability in Fig. 2. Nevertheless, the dynamics of alpha phases, as with that of the EEG topographies, also experienced some bias: the CV significantly dropped in specific time-ranges following the stimulus, to values ∼0.4 S.D. below its mean for infants and ∼1.5 S.D. below its mean for adults (Fig S6B, middle panel).

Remarkably, CV drop and VQ had partially dissociated spatiotemporal dynamics in infants. In most infant subjects, drops of alpha phase CV tended to precede VQ, as revealed by a peak at a negative latency of the cross-correlogram between the time-courses of VQ and CV (Fig. S6D). Moreover, in infants, significant VQ could be observed for longer times, even after CV was restored to baseline values due to loss of phase alignment between trials (Fig 4C). Furthermore, the spatial extension of the CV drop and VQ phenomena were generally different, as indicated by the time-courses of the numbers of channels showing significant CV drop or significant VQ (Fig S6C). In infants, the channels affected by VQ extend way beyond the range of significant CV drop (see topographies in Fig S6C). For instance, for old infants at the peak of VQ, ∼35% of EEG sites showed significant between-trial variability reduction as compared to ∼10% channels that showed CV reduction. Only for adults, the time ranges and the spatial extension of VQ and CV drop completely overlapped.

These results collectively prove the existence of a rich temporal structure in the dynamics of between-trials variability, qualitatively and quantitatively maturing over the first semester post-term birth. Furthermore, in the same infants, its temporal structure can be modulated depending on the task at hand. For infants, alpha phase reset and VQ are intertwined but distinct events: substantial VQ can exist even in the absence of an increased phase alignment between trials as made clear by analyses of variability dynamics. However, alpha phase reset might trigger VQ as, in infants, more localized and shorter-lasting CV drops precede more broadly extended and longer lasting VQ events.

### 5. Maturation of Within-trial Variability and its Relation to Alpha Phase Reset

Our third and last approach was the analysis of within-trial variability. To track within-trial variability, we quantified the amount of variation in the topography of EEG activation from one time-point to the next. This corresponds to the distance traveled in the space of possible activity topographies over a unit time or, equivalently, to the *speed* of motion in this high-dimensional space (see Methods). With this approach, topographies of activation which are stable over time and fluctuate very little from one moment to the next will yield instantaneous within-trial variability close to zero. Conversely, abrupt changes of topographies occurring at specific instants – e.g. eventual switching between microstates (Michel & Koenig, 2018; cf. Figs. 2C and S2) – would map to sudden increases of the instantaneous within-trial speed. Again, analogously to between-trial variability analyses, we related the changes of within-trial variability to the dynamics of alpha oscillations (Fig 5, S7).

**Fig 5.**
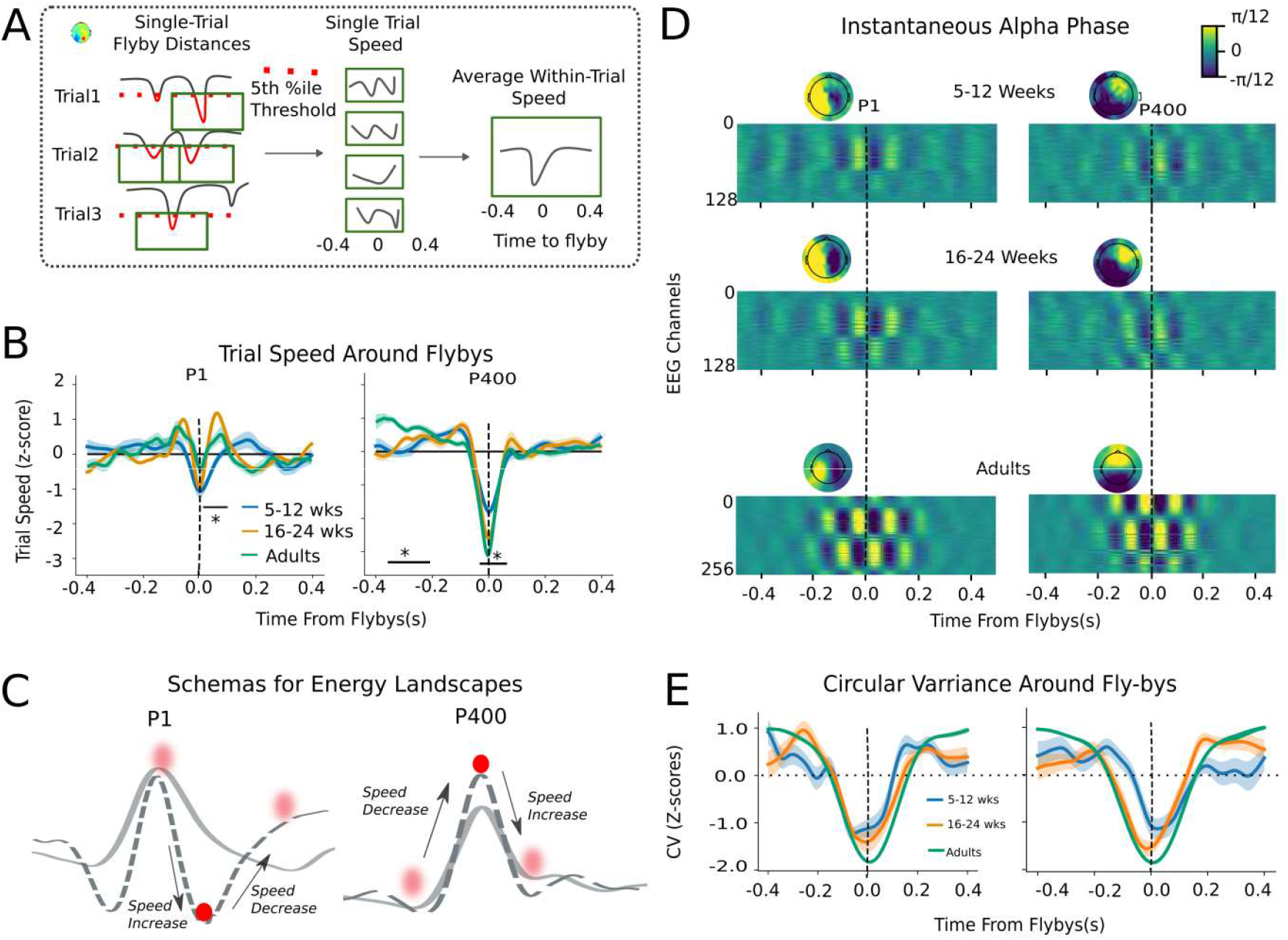
Maturation of Within-trial Speed and its relation to alpha oscillations. **A)** Trial speed around P1 and P400 flyby events for each subject was extracted by considering within-trial speed (i.e. correlation distance between topographies from one time-point and next) in 800 ms time-window around the closest 5 % flyby distances to the respective template. **B)** Group averaged trial speed profiles around P1 and P400 flybys for different age-groups. Significant speed difference existed between 5-12 weeks and 16-24 weeks old infants as indicated by black horizontal bars, cluster-based permutation t-test, p<0.05) **C)** Schematic energy landscapes that may underlie the speed profiles at P1 and P400 templates (solid line for 5-12 wo and dotted line for 16-24 wo infants respectively). The observed speed profiles could be interpreted as if motion along trajectories was driven by sampling of a structured potential energy landscape (similar to rolling of a ball across hills and valleys on a surface). **D)** Group averaged flyby triggered instantaneous phase of alpha oscillations for each channel, separated by age-groups. Y-axis represents channels ordered according to their nomenclature. Topographies were derived by plotting instantaneous phase on the scalp surface at the first peak before the closest flyby for each age group. **E)** Circular Variance (CV) of alpha phases across closest flybys. As an effect of phase-reset preceding fly-by, CV significantly reduced for all age groups ∼ [-200 200]ms surrounding the closest flyby. Shaded bars represent S.E.M. for each group.

#### Absolute within-trial variability increases with age

We first measured average within-trial variability, irrespective of the time relative to stimulus presentation. We found that time-averaged absolute within-trial variability significantly increased with age for all task conditions (Fig S7A (left panel) for lateral faces: r= 0.38, p<0.01; (right panel) for central faces: r=0.56, p<0.009). Once again, this trend continued into adulthood, with a within-trial speed (or variability) significantly higher in adults compared to infants (box plot in Fig S7A, H= 24.33, p<10^-5^, Kruskal-Wallis test). Thus, development boosted speed of exploration along neural trajectories.

We also noted, however, that within-trial speed was not homogeneous in time but had a heavy-tailed distribution of instantaneous values for both infants and adults (Fig S7 B), with extreme values possibly reflecting long jumps due to microstate switching events. Given this heterogeneity of speed modulation in time, we then moved to study whether faster or slower speeds systematically associated to specific neural configurations being visited, notably at ERP component fly-bys.

#### Speed profiles around ERP flybys are structured and modulated by age

To understand how within-trial speed is specifically modulated during the approach to known evoked ERP components, we first performed flyby-triggered averages of within trial speed of topography transitions from one moment to the next, by pulling together all individual events of closest flyby to ERP templates (possibly multiple events per trial) and averaging within-trial speed in an 800-ms window around these events (peri-flyby speed profiles, Fig 5A, also see *Methods*). Fig 5B shows average within-trial speed in the vicinity of respectively P1 and P400-flybys for lateral faces (Fig S7 C for central faces). In these profiles, peaks and troughs of within-trial speed are clearly visible, distributed symmetrically around the flyby time, and tend to get more prominent with age. Irrespective of their absolute within-trial speed, on an average, trials transiently slowed down around both P1 and P400-flybys, for all conditions and age-groups (always significantly, except for P1 flyby in adults). Slowing down was larger around P400-flyby than for P1 (condition wise signed rank test: H=30, p<0.001, H=55, p<10^-5 and H =1.0, p<0.0005 for infant central, lateral, and adult lateral faces respectively). As shown by Fig. S7 D-F (left panels), speed of exploration nearby P1 flybys increased with age, significantly for lateral faces; but not for central faces (age correlation: r=0.47, p<0.001 for left, r=0.35, p<0.02 for right and r=0.1, p>0.7 for central faces). This increase was due to the emergence of more marked positive acceleration, preceding and following the time of closest flyby. On the contrary, speed of exploration decreased with age nearby P400 flybys, due to a progressively marked deceleration, for both lateral and central faces (respectively, Fig S7 D-E right panels, age correlation: r= −0.45,p<0.001 for left, r=-0.45,p<0.004 for right and r = −0.33, p<0.07 for central faces).

These characteristic speed profiles are reminiscent of the accelerations and decelerations that a physical ball would experience when rolling on a non-flat surface, accelerating while descending into a valley and decelerating while ascending out of the valley due to gravity (see illustration in Fig. 5C). It is thus tempting to interpret these speed profiles as if reflecting the sampling of a structured landscape of effective free energy (Landau, Lifshitz and Pitaevskii, 1984). In this statistical physics-like view, neural trajectories would unroll under the influence of force fields generated by an effective energy landscape (Gu *et al*., 2018). These forces act as biases on the system’s trajectory in the proximity of critical points associated to ERP components. Through development, this energy landscape gets progressively more sculpted and the neural trajectories can be thought of as being more actively controlled. (see *Discussion*).

### ERP flybys are associated with transient phase reset but not amplitude modulation events

As in the case of between-trial variability, modulations of single-trial oscillatory dynamics may be an important contribution to the observed variations of the within-trial speed. We thus computed flyby triggered averages of the instantaneous phase of ongoing alpha oscillations at all channels (Fig 5D). In the surrounding of the closest flyby time, oscillatory patterns can be clearly distinguished, suggesting that flyby events tend to occur at similar instantaneous alpha phase. These analyses thus provide a strong evidence for phase reset in the surrounding of the flybys: (i) for all age groups; and, (ii) with a distinct global topography for each ERP component. This phase alignment between the oscillations at different ERP flybys begins ∼ 200 ms before the flyby event and ends around the same time after the flyby. These times correspond to the alpha oscillatory period with a variable decorrelation time (hence the oscillatory pattern fades away farther away from the closest flyby time in Fig. 5D,). Due to this transient phase alignment of alpha oscillations to ERP component flybys, flyby triggered Circular Variance (CV) across single-flyby events dropped significantly (Figure 5 E). However, the profile of circular variance drop was simpler than the profiles of within-trial speed variation and showed just a single minimum at flyby time, without peri-stimulus increases (unlike the speed profiles).

Importantly, within-trial speed modulations were paralleled by phase modulations but not as clearly by amplitude modulations. Considering the amplitude of the signal alpha component, alpha power surrounding ERP flybys did not significantly deviate from baseline values for any of the groups (Fig.S8A). Beyond analyses of band-restricted oscillatory dynamics, we also considered more general modulations of broadband signal-to-noise ratio, by quantifying the L2-norm of the topography of activation to track average activity levels at all channels. Averages of L2-norm of activation over stimulus-aligned trials did not show any significant upward or downward modulation in any time range following the stimulus (Fig S8B, neither for lateral faces (top panel), nor for central faces (bottom panel)). This corresponds to the fact that strong voltage activity can be found at any time within individual trials, due to ongoing fluctuations (cf. Figure 2A-C) and are not restricted to the classic ERP time-ranges. As shown by Fig. S8 C, we again found flat profiles of L2-norm change around flyby events for most combinations of age and stimulus type (the only exception being a significant increase at P400-flyby, limited to the adult group).

**To summarize,** we observed characteristic accelerations and decelerations of speed along single-trial neural trajectories in the vicinity of closest flybys to ERP components. These speed profiles were suggestive of the transiently attracting and repelling forces. They were associated with brief phase-coherent events phase-locked to ERP flybys and could be clearly detected since early infancy, despite the absence of systematic modulations of alpha or broadband power. Average speed increased with age, but transient decelerations at flyby also became more marked. Together, these findings hint at neural trajectories sampling an internal landscape of attracting configurations, with an improved sampling capacity (as the within-trial variability increases) and a more marked structuring of this landscape, as development progresses.

## DISCUSSION

In this study, we demonstrate the existence of a rich temporal organization of the neural responses to stimulus in adults and infants that goes beyond the mean response captured by ERPs. We propose the concept of Event-Related Variability (ERV) to refer to the temporally structured dynamics of the response fluctuations. To characterize ERV, we focused on the complementary aspects of: single-trial (dis-)similarity to known ERP components (ERP flyby analyses, Fig. 2); reproducibility of response trajectories across different trials (between-trials variability analyses, Fig. 3); and speed of reconfiguration of the induced activity topographies along individual trials (within-trial speed analyses, Fig. 4). We furthermore put these three aspects in relation with the dynamics of ongoing alpha oscillations to study the contribution of phase reset to ERP flybys and variability quenching. Our results confirm that, for both infants and adults, the trial variability remains very large (absolute correlation distances of order ∼0.95, hence Pearson correlation coefficient ∼ 0.05) and that stimulus presentation does not suppress ongoing fluctuations (probabilities of transition between microstates were not significantly affected). Yet this variability is significantly modulated by face presentation in a non-trivial way, as exposed by the ERP flybys and variability quenching.

In all conditions, a period of decreased variability across trial trajectories (variability quenching) is detected after stimulus presentation, confirming many previous reports in adults (Schurger *et al*., 2010, 2015; Arazi, Censor and Dinstein, 2017; Ito *et al*., 2020). Beyond these previous studies, we observe that the time-range of between-trial variability quenching evolves through age and that, at a given age, is strongly and qualitatively modified by the task at hand. Furthermore, we observe that variability quenching always occurs in time ranges in which trials approach ERP landmark topographies (ERPs flybys). The converse however is not true. This establishes ERP flybys and between-trial variability quenching as partly independent phenomena. The fact that some flybys co-occur with a between-trial variability quenching and some others not, suggest that there are different ways to approach an ERP component. Trials can approach a specific activity configuration similar to the target ERP components in a rather unconstrained fashion and hence with no change of variability between trials. On the contrary, trials can follow a specific path of approach more faithfully, producing a quenching of the between-trial variability.

Variability quenching (VQ) is also partially independent of the phase reset of ongoing alpha oscillations. The alpha phases were generally poorly synchronized across trials (circular variance between phases: ∼0.8 even at the strongest quenching). Nonetheless a slight decrease in circular variance was observed preceding VQ in infants. Importantly, variability quenching affected more extended sets of channels than the ones at which a significant phase reset was detected, especially in older infants. Furthermore, variability quenching lasted longer than the periods of phase alignment between trials following the phase reset. At the level of within-trial variability, the rich structure of within-trial speed modulations was not explained by simple variations in alpha oscillatory phase and amplitude (or by signal-to-noise ratio). Thus, reset of oscillatory dynamics may be only one facet of the complex variability modulations that system’s response trajectories experience in response to a stimulus.

### Might variability quenching denote top-down processes?

Various elements suggest that quenching events are not a mere automatic consequence of stimulus presentation, but might signal a more controlled system’s trajectory, possibly implementing a form of top-down regulation: First, quenching could occur late after the stimulus onset, for example ∼400-600 ms, i.e. in P400 time-window in older infants. Second, age did not simply extend the time-window of quenching but shifted its target, from P1 to P400, and third, even at a given age, the quenching dynamics was qualitatively modified by changes in the stimulus configuration. First-trimester infants showed P1-component flybys for both central and lateral faces, but variability quenching at P1 flyby occurred only for lateralized faces, i.e. for the most challenging task. Indeed, lateralized faces were much more difficult to perceive because they were presented briefly at a random interval and in a competition with the central attractor that helped avoid infants’ saccades. Moreover, the slow maturation of parafovea compared to fovea (Allen, Tyler and Norcia, 1996) made this brief lateral stimulus even less discernible for the younger infants. We hypothesize that variability quenching events through top-down control processes helped the infants’ guide their attention in the absence of a strong bottom-up signal. For e.g. younger infants might try to shift their attention towards the lateralized stimuli in order to “detect” it, without recognizing a specific face, or even extracting facial features, while the older infants and adults might shift their attention to the periphery only after detecting the stimulus, which then allows them to guess / recognize the face. This could explain the shift of variability quenching from the P1 component (related merely to the detection of the visual input) to the P400 ERP component that is related to the face processing. Similarly, when the face was centrally presented, younger infants could focus on the face identity, the detection part being plainly evident. From this interpretation, a quenching event (that indicates a transient constraining of the system’s dynamics to a specific trajectory) might reflect more intensive information processing than the “hit and run” visit to the P1-component observed in older participants and in the case of an easily perceived visual event (central faces) for younger participants. The existence of top-down regulatory mechanisms in infants has been confirmed experimentally (Emberson et al., 2015; Kabdebon& Dehaene-Lambertz, 2019; Kouider et al., 2015). Notably, the shift of focus for variability quenching, from P1 to P400 at ∼12 weeks (for lateral faces) corresponds to the first milestone in visual development when several peripheral structures reach maturity (e.g. lens, fovea) and myelination of the optical fibers and maturation of V1 reach a plateau after a period of rapid change. It translates in the convergence to adult values of the P1 latency for centrally presented stimuli around 12 weeks (Dubois *et al*., 2008), while peripheral vision matures more slowly(Allen, Tyler and Norcia, 1996). Feed-back connectivity is also progressively restructured after term-birth, passing from disperse growth to selective pruning (Kennedy *et al*., 2007) which may allow for more effective attention control or predictive influences.

We note, finally, that, in adults, modulations of alpha oscillatory activity have been interpreted too as signatures of top-down control and attention mechanisms (Jensen, Bonnefond and VanRullen, 2012; Klimesch, 2012), although with some controversy on the relative contribution of power and phase changes (Van Diepen *et al*., 2015). Here we found the evidence for phase reset preceding ERP component flybys but not for consistent power changes (Fig. S6 B and S8 A). Modulations of alpha oscillatory dynamics and more general reductions of trajectory variability (robustly detected at all ages) co-occurred in adults suggesting a potential equivalence (Daniel *et al*., 2019). As we have seen, however, in infants, the two phenomena are partly decoupled (cf. again Figs. 4 C and S6 C). Thus, the study of an early infancy developmental window provides a unique opportunity to disentangle two possible mechanisms that could underlie attention modulations: on one side, timed and selective inhibition, provided by alpha oscillations reconfiguration (Foxe and Snyder, 2011); and, on the other, reduced noise (Arazi, Yeshurun and Dinstein, 2019) and controlled selection of state-specific trajectories (Baria, Maniscalco and He, 2017; He, 2018). In future, task difficulty and information processing load should be parametrically adjusted to investigate their respective influence on ERV dynamics and its coupling to modulations of alpha oscillatory activity, from early infancy to adulthood.

#### Within-Trial Variability Modulation

Beyond the common focus on modulations of variability between trials, we emphasized variability modulation events taking place within individual trials. We confirmed previous results obtained for adults (Schurger *et al*., 2015) and extended them to infants. If a between-trial variability quenching event denotes that the flow of system’s trajectories is restrained to a specific manifold when reaching and leaving an ERP component, the phenomenon of within-trial slowing-down suggests that the flow of *each of the individual trajectories* on this manifold characteristically decelerates when the system approaches certain landmarks (ERP component flyby). This is important, because perception and cognition happen in real time (without waiting for multiple stimulus presentations before perceiving a face). Therefore, instantaneous modulations of response variability can be instrumental only if they occur within individual trials. The slight slowing-down of individual trials near P1- and, in a particularly marked way, P400-flybys, correspond to the system trajectories lingering in an orbit around the corresponding ERP template for a short time. Such transient restriction in the system’s fluctuations –or even the boosted speed when entering or leaving the orbit, as for P1 flybys might be detected, e.g., by an integrator neuron –serving as a tempotron readout (Gütig&Sompolinsky, 2006)– to signal that a given stage in cognitive processing has been reached and thus initiate the next processing step in a sequence (Zylberberg *et al*., 2011). These speed modulation profiles can remain well identifiable by the system, despite the large variability of spatial topographies at ERP flybys.

Combined together, our results suggest that more than the current position of the system in its configuration space, ERPs are marked by “how” the system is flowing through and away from its current position. The evolution of the system seems far from being at a stable attractor. The fact that dynamics is dominated by structured fluctuations make such a system compliant with reservoir computing systems (Maass, Natschläger and Markram, 2002), in which intrinsic chaotic fluctuations are only transiently reduced by the applied inputs and are actually needed to boost learning capabilities.

#### Mechanisms underlying variability modulations and their possible functional relevance

The question remains as to what mechanisms could be responsible for the emergence of such a structured ERV components. This question has been explored in some depth for the quenching of firing rate variability in neuronal population responses to a stimulus (Churchland, Yu, *et al*., 2010; Fairhall, 2019), where mechanisms such as attractor stabilization (Litwin-Kumar and Doiron, 2012), chaos suppression (Rajan, Abbott and Sompolinsky, 2010) or “supra-linear stabilization” (Hennequin *et al*., 2018) have been proposed. Large-scale computational efforts have demonstrated that such microscopic properties could indeed be an ingredient for the macroscopic variability quenching(Ponce-Alvarez *et al*., 2015) which are sampling a more global activity than the neural firing recordings reviewed by Churchland *et al*. Contrarily, some studies have suggested that macroscopic quenching could be due to the variations in the baseline and/or the phase of ongoing alpha oscillations (Dinstein, Heeger and Behrmann, 2015; Daniel *et al*., 2019). These views are akin in spirit to earlier and more recent proposals that oscillatory phase and amplitude modulations are responsible for the generation of ERP components and their trial-to-trial fluctuations (Hanslmayr *et al*., 2007). Here we have found indeed that phase reset events tend to precede the detected between-trial variability quenching events (Fig. 4C, Fig S6). However, variability quenching can persist longer even after phase reset and the return to baseline levels of circular variance of alpha phases across trials. This means that neural response trajectories keep being restrained to a common manifold even when ongoing oscillations are not anymore aligned in phase, due to spontaneous decorrelation. A more general form of trajectory control may thus be acting, and alpha phase reset could be an early component of the implied control mechanisms, if not their causal trigger (as phase reset seems to precede quenching), or, alternatively, just an epiphenomenal manifestation of them.

Moreover, the characteristic kinematics of system’s evolution was observed here as neural trajectories approach ERP components. Such type of kinematics and effective landscapes may naturally emerge because of the non-linear dynamics of multi-scale neural circuits, self-organizing into structured flows on low-dimensional manifolds (Pillai and Jirsa, 2017). The occurrence of coherent oscillations and phase reset dynamics at ERP flyby (Fig 5D) would not be incompatible with a structured internal landscape of the attracting states being sampled. Indeed, nonlinear oscillators can respond with endogenously-generated phase shifts to external forces (Acebrón *et al*., 2005; Kirst, Timme and Battaglia, 2016), and the effects of approaching a dynamical critical point at ERP flyby can be conceptualized as forces acting on the (oscillating) system’s trajectory. Thus, phase reset events could be precisely caused by the curvature of the internal effective energy landscape being sampled. Note that, if the observed speed profiles had to be explained as due to motion in a force field, the free energy minimum of this landscape would then be located at the first maximum of speed, which precedes the time of closest passage near ERP templates by ∼50-100ms. Thus, ERP-like configurations, associated to the lowest speed, would not be the “attractors” but rather signal the moment of crossing from one critical point to the next.

In this framework, changes of ERV dynamics through development and learning would be accounted for by the growth of more marked barriers and sinks in the effective energy landscape or, equivalently, bifurcations causing the birth, fusion or death of different attractors or saddle points in the system’s high dimensional phase space. Such conjectures may be potentially validated by estimating the morphology of an effective free energy landscape surrounding the ERP templates (Ezaki *et al*., 2017). Similar to previous studies, we find here also the increase of overall within-trial variability along early development (Fig. S7 A-B) (McIntosh *et al*., 2010; Garrett *et al*., 2011). This increased structuring of the landscapes that shaped the brain activity may mediate the capability to learn internal statistical models of environment, for better inferences in perception (Berkes *et al*., 2011).

Additional theoretical and computational investigation will be needed to disambiguate which of the possible scenarios is leading to the observed quenching. Indeed, different models may predict different statistical distributions of secondary features –such as e.g. the jitter in latencies from stimulus presentation (or phase reset events) to ERP component flybys– to compare with the empirically measured ones. Furthermore, parameters such as the local excitation/inhibition balance within cortical populations evolve with age (Hensch&Fagiolini, 2005) and the maturation of these parameters may predict different ERV developmental trajectories depending on the actual dynamic mechanism. To probe the functional relevance of ERV, a focus on an early developmental period (and, particularly, as early as the first trimester of life), may be once again crucial. Indeed, learning within functional architectures not yet fully scaffold is exceptionally fast (Dehaene-Lambertz and Spelke, 2015). At the same time, ERV dynamics is already rich and still evaluative and separable from phenomena such as alpha modulations that are dominating in adults and may be concealing subtler functional aspects of response variability.

#### Methodological considerations

Our approach has methodological limitations that could be overcome by future developments. For instance, the extraction of ERP templates in our case depended on a manual inspection based on the developmental literature, but more sophisticated algorithms for the temporal clustering of single-trial ERP topographies (Vahid *et al*., 2020) could be used and combined with our variability analyses schemes. The occurrence of large within-trial variations and, particularly, of extreme within-trial speed values at certain times (Fig. S7 B)– also need to be put in correspondence with the scale-free dynamics of microstate transitions (Van de Ville, Britz and Michel, 2010) or other approaches to describe spontaneous dynamics as random walks in high-dimensional state spaces, which also correlate speed variations with development and cognitive performance and are shaped by intrinsic dynamical landscapes (Hansen *et al*., 2015; Battaglia *et al*., 2020; Lombardo *et al*., 2020). Finally, linking activity to network state dynamics –what is the functional connectivity triggered by an ERP flyby?– may allow assessing whether exchange of information is dominated by bottom-up or top-down flows at different ages or ERP stages (Bastos *et al*., 2015). Note that our measures were very sensitive to age, allowing capturing even subtle differences between the left and right hemisphere maturational calendar. The ERV approach might thus be a more sensitive tool than classical ERPs to capture differences between experimental conditions in infant cognitive studies, but above all to explore neurodevelopmental disorders.

#### Conclusion

ERPs have been an attractive description of the post-stimulus brain activity, described as successive steps defined by their reproducible latency and brain sources, allowing obtaining neural algorithms underlying cognition. However, this description was somewhat misleading, ignoring the ongoing activity. The framework proposed here encompasses both aspects. We recovered that ERP components serve as a “compass” to identify special dynamical points for the on-going activity sampling erratically vast volumes of the neural configurations space, confirming that ERPs are indeed capturing neural consequences of a stimulus presentation. At the same time, we also showed that they are far from capturing the entire activity patterns following a stimulus. We proposed the term ERV, as a better concept to describe the neural consequences of a stimulus. This proposal is not purely semantic, since it allows describing the ERP maturation in integrated manner on one hand and emphasizes the structured variability of the background EEG on the other. It allows thus speculating that the gradual change of ongoing activity might reflect the increasing knowledge of the environment throughout development of a structured internal landscape biasing neural trajectories (that, on their turn, through their volatility, can efficiently sample this landscape).This approach might be particularly fruitful to investigate neurodevelopmental disorders and their cognitive consequences.

## MATERIALS AND METHODS

### Subjects

Reported results included data from two cohorts. The first group of healthy full-term infants (N = 39, Mean age: 14.15 ± 4.79 weeks, age range: 5.6 to 23.6 weeks, 11 girls) was studied elsewhere to investigate the functional maturation of visual Event Related Potentials (ERP) to lateralized faces (Adibpour, Dubois and Dehaene-Lambertz, 2018). A subset of these infants (N = 22, Mean age: 14 ± 4.96 weeks, age range: 5.6-22 weeks, 7 girls) was also tested to study ERP responses to central faces. To compare the results obtained for infants with adults, we additionally included a second group of young adults (N= 13, Mean age: 23.39 ± 2.32 years, age range: 21 to 27.1 years, 6 females) who were presented with the lateralized faces following the same paradigm as infants. The study was approved by the ethical committee for biomedical research. All adult subjects and parents of infants gave written informed consents before participating in the study.

### Task Paradigm and Protocol

#### Lateralized Faces

Each trial started by a rotating colored bull’s-eye that remained at the center of the screen during the whole experiment to attract infants’ attention to the center of the screen. Streams of face images (male or female face out of 6 neutral, unfamiliar front faces) appeared consecutively on the left and right side of the bull’s eye for 250 ms followed by a variable delay between images (550 to 950 ms post-offset of the image with a 50-ms step). The asynchronous presentation ensured minimal anticipatory gaze to the left or right side. To investigate the inter-hemispheric transfer of information in infants, each stream included three types of images: a side-assigned face image (standard), a novel face (new-deviant), or the face commonly assigned to the other side (known-deviant), with the expectations that an efficient inter-hemispheric transfer ensures ERP response to known-deviant faces to be similar to standard faces. Each block included ∼80 % standard, ∼10 % new-deviant and ∼10 % known-deviant faces. For the current analyses however, we considered all faces presented on either left or right side; irrespective of this distinction.

#### Central Faces

One female and one male face, not used during the lateralized paradigm, were presented at the center of the screen for 250 ms, spaced by a random interval of 550-950 ms during which the colored bull’s eye was presented.

#### EEG Protocol and Pre-Processing

EEG recordings were acquired with EGI net comprising 128 electrodes for infants and 256 electrodes for adults, and digitized in real-time at a sampling rate of 250 Hz. EEG data was further pre-processed in EEGLAB software. Recordings were band-pass filtered between 0.5 and 20 Hz, the signal was segmented into epochs of 1.9 s (−0.4 to 1.5s relative to the onset of face presentation). Channels and trials contaminated by motion or eye-blink artifacts were rejected. For infants, additional trials were rejected when the eye-gaze moved away from the screen; by manual inspection of video-recordings. Epochs were re-referenced by reference averaging but no baseline correction was applied to allow unbiased analyses of post-stimulus variability as compared to pre-stimulus variability. Finally, EEG topographies were normalized by dividing the activity of each sensor by the global field power (GFP, i.e. standard deviation across sensors) at each time-point. For the current analysis, further temporal smoothing was applied by averaging the activity at each sensor in a 100-ms overlapping sliding window centered at a given time point in each trial (all results were validated without this temporal smoothing). Additional information about data acquisition, pre-processing and task paradigm not pertaining to the current study is detailed elsewhere (Adibpour, Dubois and Dehaene-Lambertz, 2018). For infants, final dataset considered for further analyses included ∼110 ± 60 trials (min= 38, max = 246 trials) each for left and right faces, and ∼ 32 ± 17 trials (min= 3, max=74 trials) for central faces condition. For adults, the final dataset included ∼353 ± 33 (min= 255, max=363 trials) trials each for the left and right faces.

### Trajectory in Principle Component (PC) Space

For one example subject, 12s long segment of clean continuous EEG data containing 10 consecutive (left and right faces) trials were low-pass filtered using 100 ms overlapping sliding window. This segment of EEG data was normalized by dividing the activity at each sensor by the instantaneous global field power. This standardized 128-dimensional time *x* channel matrix was transformed into three orthogonal components that explain a maximum amount of the variance (82% of total temporal variance was explained by first 3 components) using PCA decomposition from scikit-learn toolbox in python and the resulting PC coefficients were used to visualize 3-dimensional trajectory shown in Fig 2B

### Microstates Analysis

#### K-Means Clustering

To derive combined microstates for infants, 128-dimensional continuous EEG signals were concatenated across time dimension for all infants after the preprocessing steps were performed. Bad segments were ignored from the analysis and bad channels were interpolated using linear spatial interpolation. Each individual topographic pattern at time t was normalized by GFP before passing it to the further analysis. Python library scikit-learn was used for clustering continuous data into discrete microstates, with number of clusters pre-defined (n=4). The 4 cluster means thus identified were than considered as “microstates” (shown in Fig 2C top panel). Nearest neighbor algorithm was then used to assign microstate labels to each instantaneous topographic pattern. That is, for each subject, the instantaneous topography at each time-point was compared to the 4 microstates using correlation distance (1-Pearson correlation) and a microstate closest to the instantaneous topography at time t was assigned as a label at this time. For each subject, the symbolic sequences of microstate transitions (labeled from A-D) were further segmented into epochs to align them to stimulus onset times.

#### Microstate Transition Trains and Transition Probabilities

Each epoch of labeled sequences was then binarized to encode a transition: i.e. if the microstate changed from the current one to the next at time t, it was encoded as a “spike” (or 1) at the time t. These transition trains (visible in Fig 2C) were further smoothened by convolving them with a 100ms smooth Gaussian kernel. These smooth transition curves were averaged across trials to convert the transition trains into transition probabilities (Fig 2C Bottom panel).

#### Microstate Observation Probability

To calculate probability of observing a specific Microstate at time t after stimulus presentation, the epochs of labeled microstate data was simply one-hot encoded and summed across trials for each microstate individually in time-aligned manner. These observation probabilities were then averaged across subjects for comparison across age-groups (Fig S2).

### Extracting ERP Templates

For each condition (left, right and central) and for each cohort (infants and adults), we derived grand average ERP topography by averaging subject-specific ERP activity separately for each sensor across subjects (Fig 1B for infants, Fig S1A for adults). For infants, we identified ‘P1 template’ as grand-average topography in the range of ∼225-275 ms post-stimulus for lateralized faces and in the range of ∼125-175 ms post-stimulus for central faces. Similarly, ‘P400-template’ was derived as the average topography in the range of ∼525-575 ms post-stimulus for both lateral and central faces (Fig S3 A). For adults, we identified ‘P1 template’ as the grand average topography in ∼75-125 ms post-stimulus while ‘P400 template’ was identified as ∼375-425 ms post-stimulus (Fig S3 B). These time-ranges were chosen by selecting a 50ms long time-window around the peak ERP response topography as inspected manually.

### Measures of Trial-Variability

Measures of trial-variability (i.e. flyby to known ERP templates, between-trial variability and within-trial speed) were calculated as topographic dissimilarity using spatial correlation distance (1-Pearson correlation coefficient) as dispersion metric. Hence, absolute distances varied from 0 (absolute correlation) to 2 (no correlation). Correlation distance decouples the topographic patterns from their magnitudes, allowing focusing on the relative spatial patterns rather than their absolute magnitudes. Note that for our study, correlation distance was mathematically equivalent to previously used cosine dissimilarity measure since our data was reference averaged at each time-point (hence, mean across sensors equals to zero).

### ‘Flyby’ to ERP Templates

For each subject and for each condition, flyby distance from trial *κ*to a certain ERP template *Χ* at time *τ* was calculated as correlation distance,

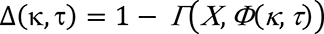

Where, Γ is Pearson’s correlation coefficient and *Φ*(*κ*, *τ*) represents topography at trial *κ* and time *τ*. These single-trial distance time-series were further averaged across trials for each ERP template to obtain a single time-series per subject for P1 and P400 templates and for each condition (Fig 3 A).

### Between-Trial Variability

For each subject and for each condition, between-trial variability at time τwas calculated as the average of all pair-wise spatial correlation distances between all trial-pairs *i* and j;

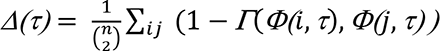

Where, *n* = number of trials, 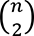 suggests all pair-wise combinations of trials, *Γ* is Pearson correlation coefficient and *Φ(j*, *τ)* represents sensor topography at trial *j* and timepoint*τ*.

One such absolute single-subject between-trial variability time-courses were derived per condition and further z-scored across time, to obtain relative between-trial variability. These z-scored time-series were further averaged in the previously defined time-range for P1 and P400-flybys to obtain relative between-trial variability around flybys (Fig 4, Fig S5).

### Topography of Between-trial Variability Quenching

If sensor χ has δ neighboring channels, Between-trial variability is calculated for this sensor at time τ as follows:

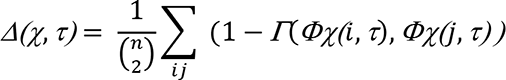

Where, *Γ* is Pearson correlation coefficient; (*Φχ(i*, *τ*) is *δ* + 1 dimensional activity vector at time *τ* and trial *i*, where each dimension represents neighbors of sensor *χ* (including itself). Neighbors of each channel were inferred from the channel-connectivity matrix estimated using find_ch_connectivity function of MNE-python. For each subject, instead of single global between-trial variability time-series, now we obtained one time-series each for each sensor. This absolute sensor-level between-trial variability was further Z-scored across time to obtain relative variability for each sensor. These channel x time matrices for each subject were further averaged to obtain group-level between-trial variability topography (Fig S6).

### Quantifying Features of Ongoing Alpha Oscillations

For each subject and conditions, pre-processed epochs of EEG signals were first band-pass filtered in a narrow frequency band of 9-12 Hz using MNE Python’s default FIR filter. Analytic signal ( *y*_n_) were than derived for each channel as follows:

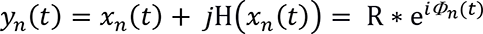

Where Η(*x*_n_(*t*)) represents Hilbert transform of the original signal*x*_n_(*t*). This analytical signal discards negative frequency components without loss of information and makes instantaneous phase *Φ*_n_(*t*) of the signal accessible. Moreover, Circular Variance (CV) i.e. the variability in instantaneous phases (or phase asynchrony) can be calculated simply as follows:

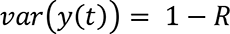

Where, *R* is the Kuramoto Order Parameter that determines the average phase synchrony across trials.

The CV varies between 0 to 1 with 0 suggesting complete s and 1 suggesting that all trials have completely misaligned phases at time *t*.

Moreover, to assess instantaneous amplitude envelope or alpha power at each time-point, we simply consider the absolute value of analytical signal, i.e.|*y*_n_(*t*)|. These quantities were derived for each channel*n*, at each timepoints *t*and averaged across channels to evaluate group-level differences in Fig S6 for between-trial CV and amplitude as well as between flyby snippets in Fig 5 and S8 for evaluating phase synchrony and alpha power around flyby.

### Within-Trial Speed

For each trial k, within-trial speed at time *t* was calculated as spatial correlation distance between topography at that time-point and the same at the consecutive time-point.

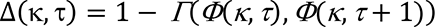

Where, *Γ* is Pearson correlation coefficient and *Φ*(*κ*, *τ*)represents sensor topography at trial *κ* and time-point *τ*. Overall absolute within-trial speed for each subject was obtained by first averaging speed time-courses across trials and then averaging the mean speed time-courses across time (Fig S7A).

#### Fly-by Triggered Speed Profiles

For each subject, moments of ‘fly-by’ to known ERP templates were identified as described above (i.e., trial-to-template distance falling in the lowest 5thpercentile). Trial-speed segments at each occurrence of ‘fly-by’ were extracted as speed time-course from 400-ms before to 400-ms after ‘fly-by’. Each of these 800-ms long speed time-courses were averaged to obtain a single speed profile per template (Fig 5A). These subject-specific absolute speed profiles were z-scored along the time dimension to obtain relative speed profiles around fly-by and further averaged to obtain group-average speed-profiles (Fig 5B; Fig S7C). To compare flyby triggered instantaneous speed across infants, peak speed (for P1 template) and lowest speed (for P400 template) were identified in the 100 ms time-window centered at ‘flyby’ moment (Fig S7D-F).

#### Trial Speed Distribution

To obtain trial-speed distribution for each age-group per condition, all single-trial speed time-courses were concatenated along time and along subjects in that age-group to obtain one single speed distribution. Probability density was obtained by normalizing area under each bin to 1. Normalized bin-counts (density) were plotted on a log-log scale (Fig S7B).

#### Fitting power-law to the Trial Speed Distributions

We first temporally concatenated all the trials for each subject per each condition and age-group cohorts. This allowed us to reliably estimate the parameters for heavy-tailed distributions. We transformed the distribution of trial-speed into the standard normal distribution and finally fitted a least square regression line to the section of log-log plot achieved from the standard normal-distribution. We further repeated the procedure for each subject and obtained slopes and biases of the best-fit lines per subject and compared across the age-groups.

### Statistics

Potential linear age-trends were tested using one-tailed permutation test on Pearson Correlation Coefficient (number of permutation=1000). Significant reductions in variability time-courses were tested using one-sample t-test and correction for multiple comparison and temporal non-independence was applied using cluster based permutation test as implemented in MNE-python(Gramfort*et al*., 2013). Group-level differences between paired groups of variables (variability in lateral vs central faces) were tested using nonparametric two-tailed Wilcoxon signed-rank tests (from Sci-py package). Group differences between 5-12 week-old (first trimester) infants, 16-24 week-old (second trimester) infants and adults were tested using non-parametric Kruskal-wallis test (Sci-pyimplementation), followed by post-hoc pair-wise comparisons using Mann-Whitney U test with Bonferroni correction (using scikit-posthocs package).

## Notes

### Competing Interest Statement

The authors have declared no competing interest.

### Summary of Updates

The manuscript was extensively revised to link the current work with resting state oscillatory dynamics.

