## Supplementary Figures for "Event-Related Variability is Modulated by Task and Development"

**This file contains:**

Supplementary Figures S1 to S8 and captions

**Note:** All faces presented as stimuli are replaced by gray squares in adherence to the Biorxiv privacy policy. The squares are representative of the relative position where originally the face stimuli appeared.

### Supplementary Figures

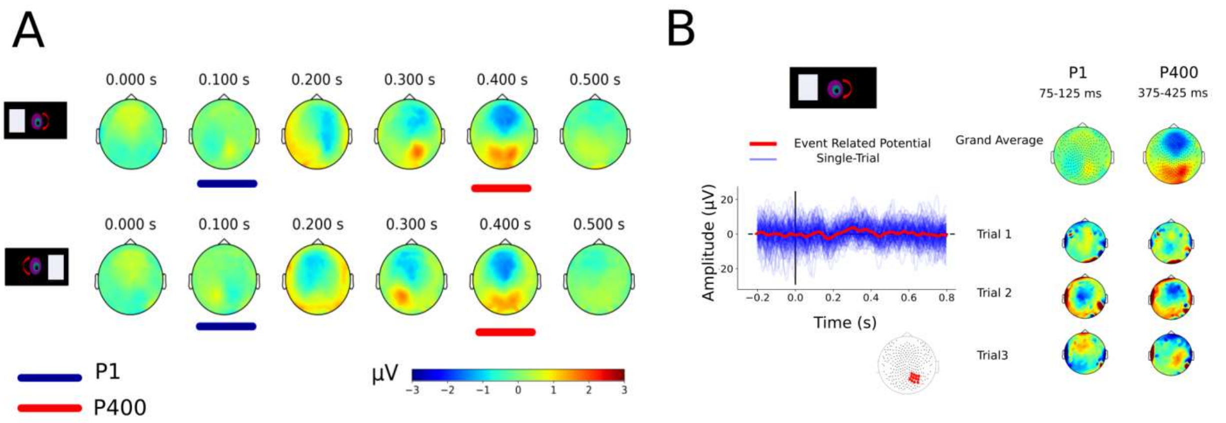


**Fig S1 Variability of Adult ERP Responses.**

**A)** Grand average spatiotemporal responses for adults when they were presented with faces consecutively on the left and right hemi-field (top and bottom panel respectively). Voltage topography of early (~ 100 ms) and late (~400 ms) ERP components were similar to those observed for infants. Hence we considered these topographies as P1 and P400 response topographies for adults (marked in blue and red respectively). **B) (Left panel):** Example voltage time-courses averaged across contra-lateral occipital electrodes for one representative adult for faces presented in the left hemi-field. Average (ERP) time-course (red) is notably weak as compared to strong single-trial fluctuations (blue). (**Right panel):** Single-trial topographies of P1 and P400 responses are notably very different from the grand average as shown for the example trials.

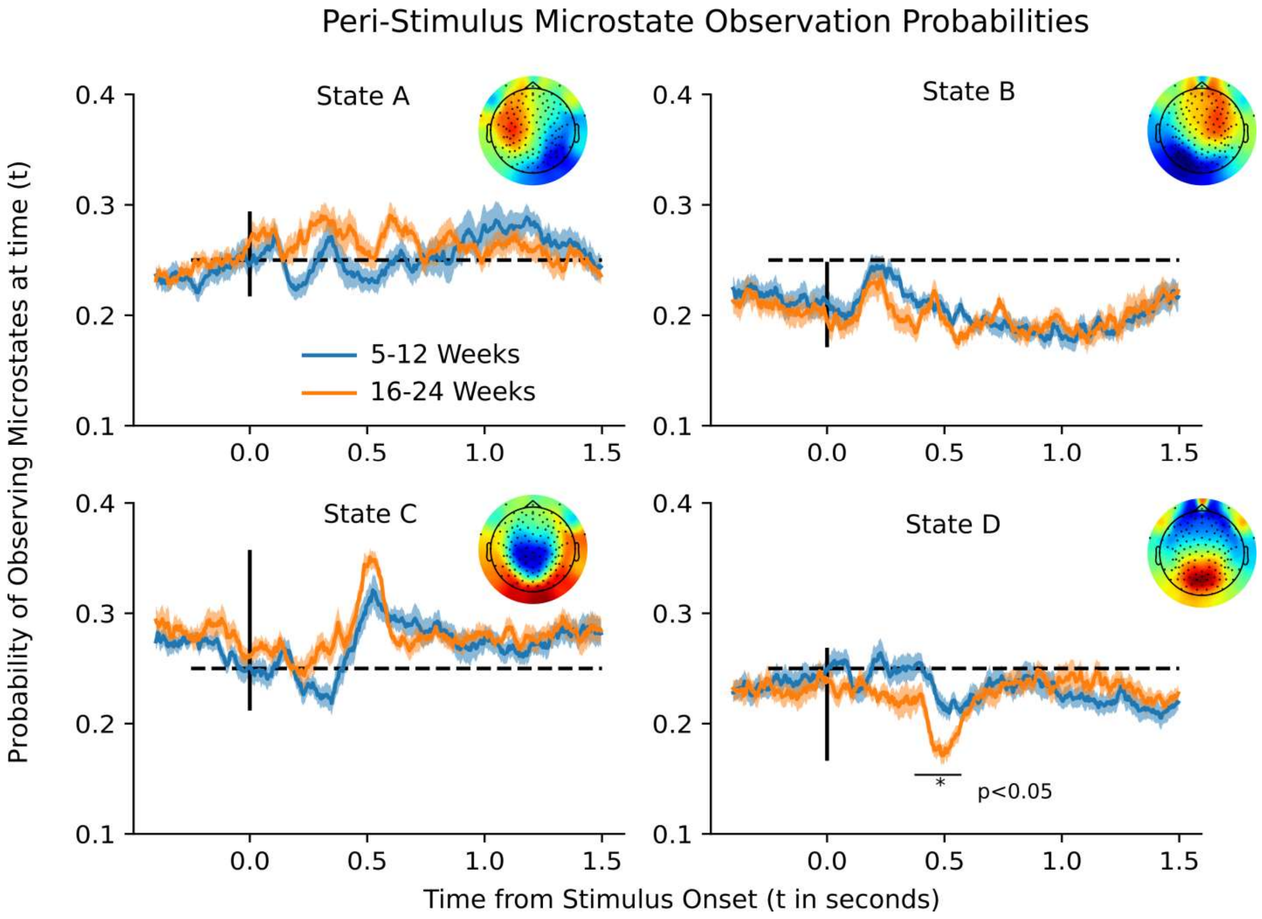


**Fig S2 Microstate Modulations in Peri-stimulus Duration. A-D)** Probability of observing a specific microstate at time (t) after stimulus onset (vertical black lines), averaged across trials and across participants separately for the young (5-12 weeks) and old (16-24 weeks) infants. For all microstates, this probability remained at chance level (~ p(obs) = 0.25, indicated with the black horizontal dashed line) at all times. ~500 ms post stimulus probability of observing microstates C (and D) were significantly above (or below) chance level, with the significant age difference observed for the state D. Shaded regions indicate S.E.M. Microstate topographies are shown on the right of each panel for the reference.

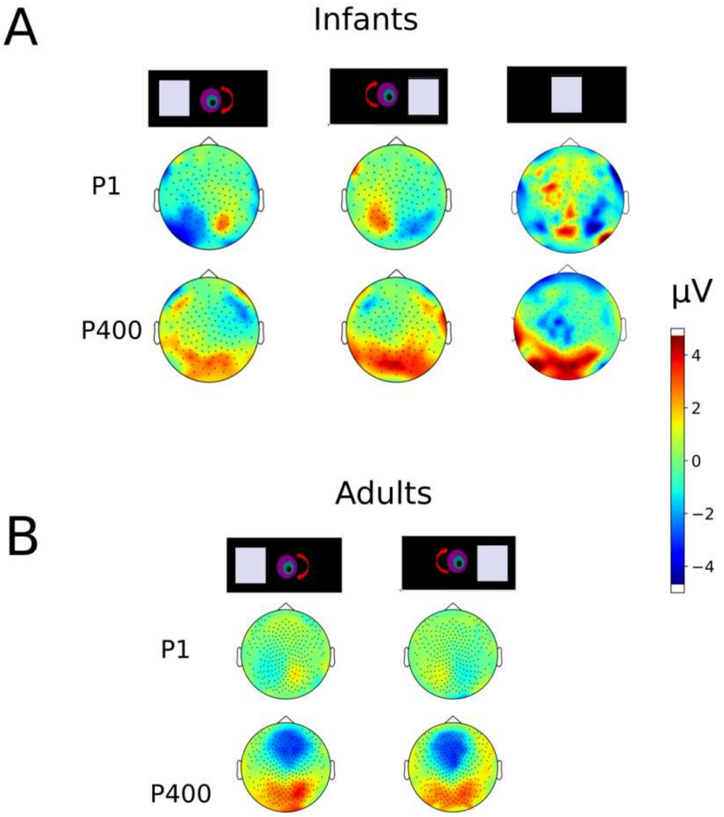


**Fig S3 Grand Average ERP Templates** for the early (P1) and Late (P400) ERP components in each condition (lateral or central stimulation) for infants **(A)** and for adults **(B).** For infants we identified "P1 template" as average topography in the range of ~225-275 ms post-stimulus for lateralized faces and in the range of ~125-175 ms post-stimulus for central faces, "P400-template" was identified as the average topography in the range of ~525-575 ms post-stimulus for both lateral and central faces. For adults, we identified "P1 template" as the grand average topography in ~75-125 ms post-stimulus while "P400 template" was identified as ~375-425 ms post-stimulus. The selection of time-windows was based on 50ms window surrounding the peak response as derived from visual inspection of topography time-series.

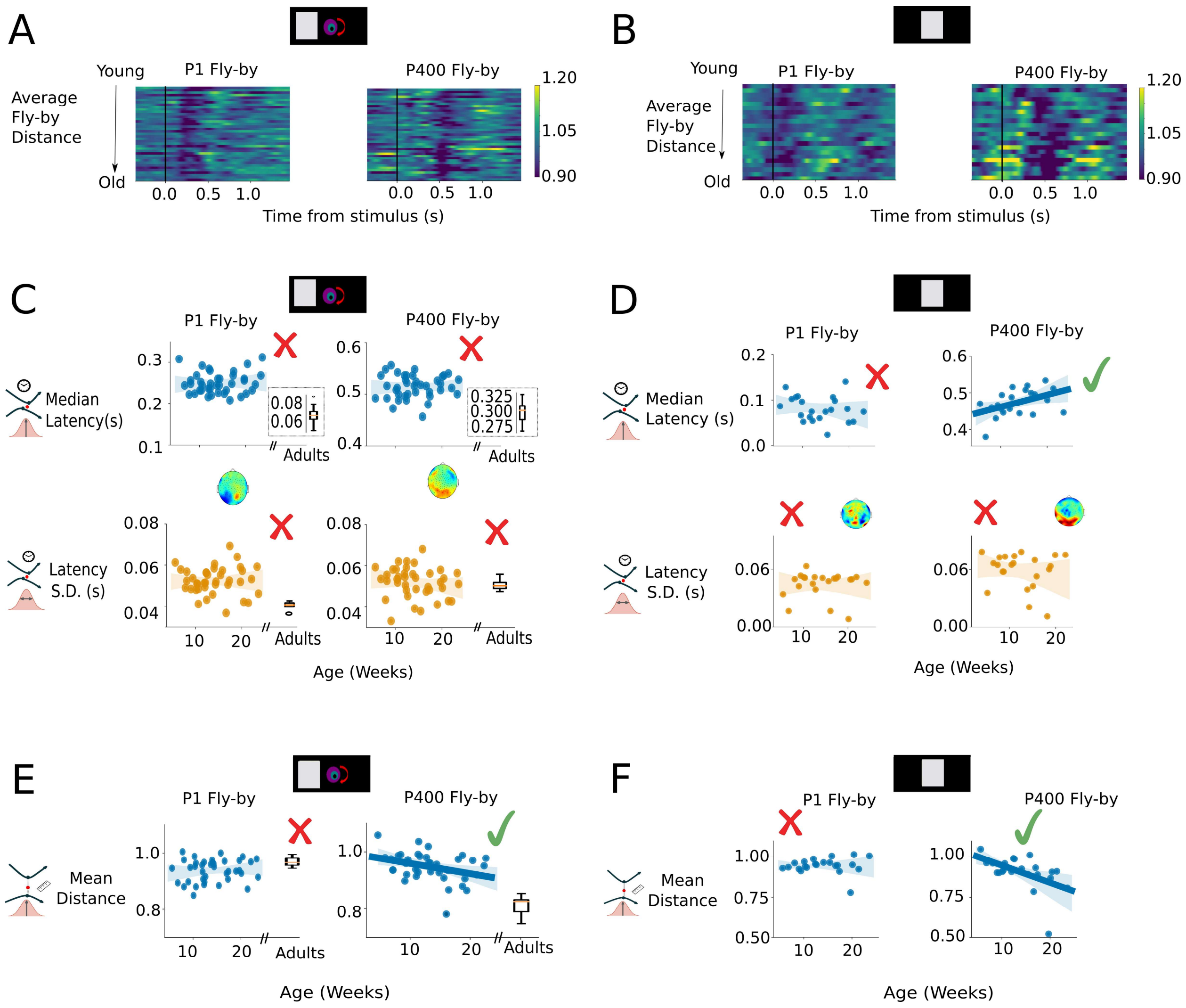


**Fig S4 Differential Maturation of Single-trial Fly-bys for Different Conditions.**

**A-B)** Average fly-by distances to P1 and P400-like templates for each infant. Each row represents a single infant, sorted in ascending order according to their age. (**A**) For left faces, trials on an average passed closer in the range of [150,350] ms for P1 and [400,600] ms for P400 template and (**B)** For central faces in the range of [-150, 150] ms for P1 and [350,550] ms for P400 template. **(C & D)** No significant age trend was observed, neither in median (top) nor in S.D. of the latencies for left faces ($r<0.01$, $p>0.2$ for all correlations) or central faces. Except for the P400 median latency ($r= 0.46$,$p<0.02$ ) which was related to increase in ERP latency itself.

Inset box-plots represent this statistic for adults when available. **(E & F)** For both conditions**,** mean fly-by distance amplitude to P400 template (top right panels) significantly reduced with age ($r = -0.35$, $p<0.018$ for left faces, $r = -0.52$, $p<0.006$ for central faces) but not for P1 template (top left panels) . r-values corrected for multiple comparison with one-tailed permutation test. Box plots represent statistics for adults. When significant, the age trend were extended until the adulthood. (Non-)Significance of linear regression is marked by (red or) green checkmarks.

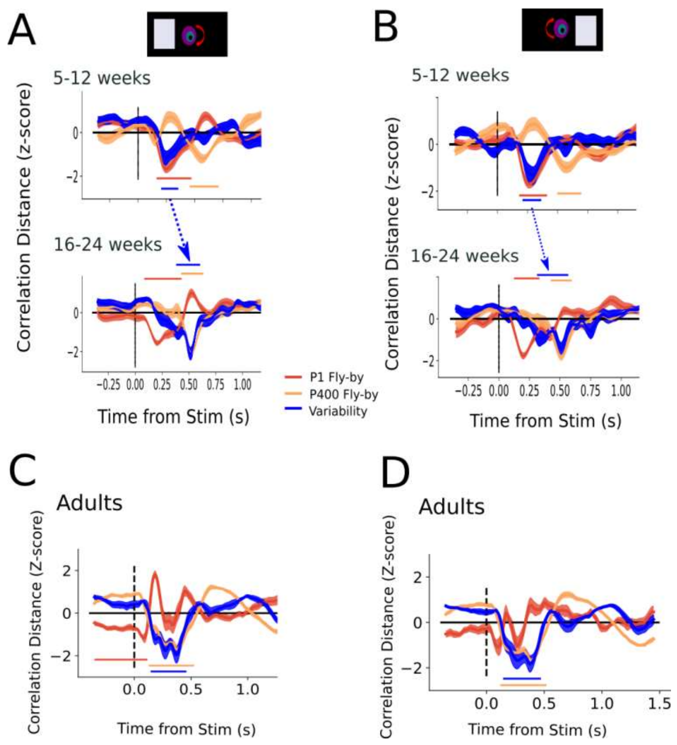


**Fig S5 Condition Wise between-trial Variability for lateral faces** for infants (A & B) and adults (C & D). Blue lines indicate group-averaged between-trial variability z-scored across time. Red and orange lines indicate group averaged fly-by distance from their corresponding P1 and P400 templates respectively. Horizontal lines indicate significant reductions from average. (One-sample t-test, cluster based permutation test, $p<0.05$). For both left and right faces, young infants reduced variability around P1-flyby while old infants and adults reduced variability around P400-flybys. Since there was no qualitative difference between the two lateralized faces, we chose to combine them.

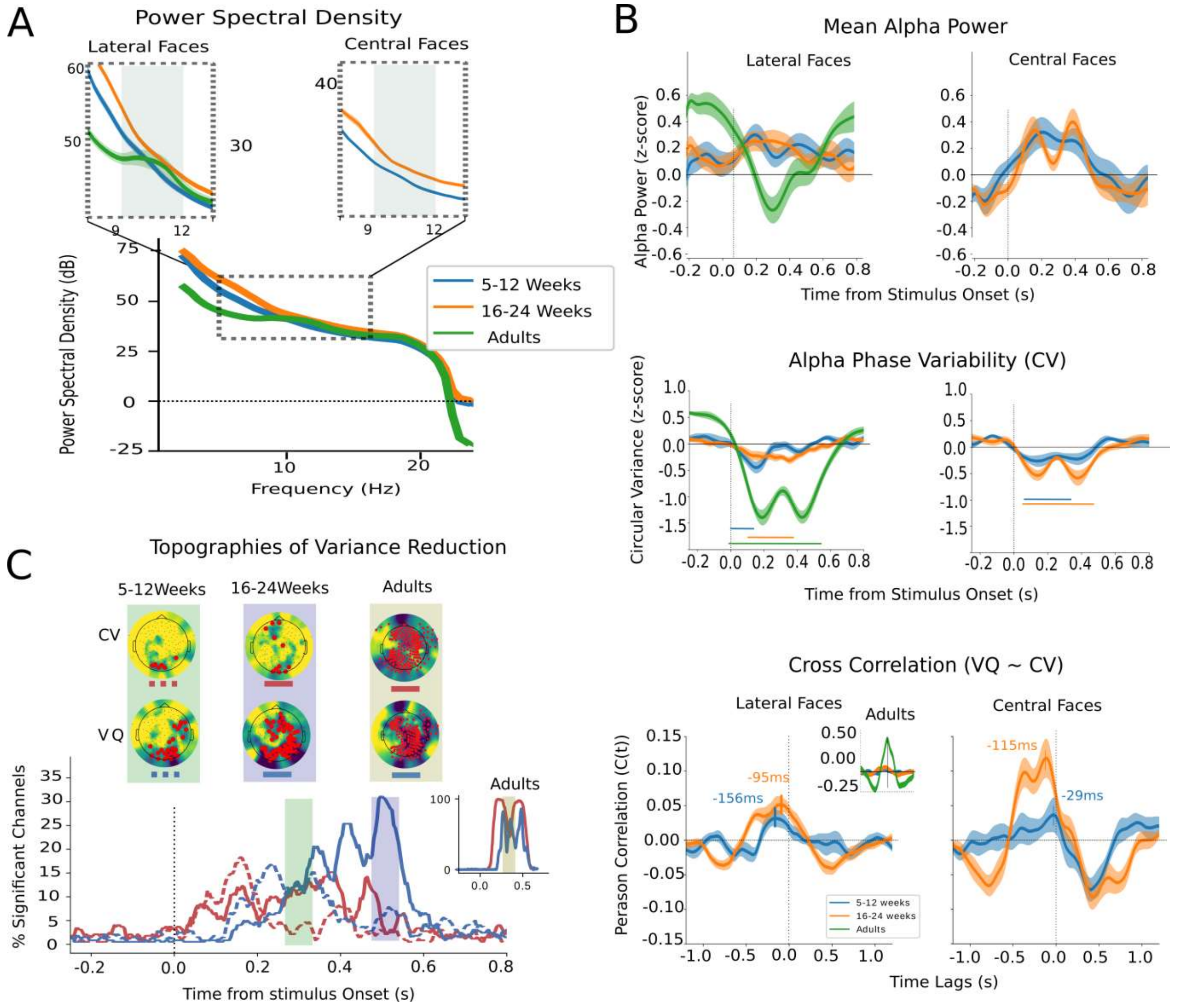


**Fig S6 Relationship between Variability Quenching (VQ) and Ongoing Alpha Oscillations.**

A) Grand averaged power spectral density (PSD) curves averaged across all electrodes, epochs and subjects for each age-group and condition separately. No prominent peak observed for infants, while for adults power increase in 9-12 Hz range is observed (visible as a bump in the green curve in the zoomed inset). B) (top panel) Channel-averaged, stimulus-aligned amplitude envelope of EEG signal band-pass filtered in alpha frequency and averaged across subjects for each age-group and for lateral and central faces (in left and right panel respectively). (Bottom panel) Circular Variance (CV) of instantaneous alpha phases across trials for the same signals.CV reduced significantly for each groups in specific time-range after stimulus presentation ($p<0.05$). (C) Time-series of Percentage of EEG channels with significant reductions in Between-trial variability or VQ (in blue) and reductions in CV (in red) for the young (dashed lines) and old (solid lines) infants and for adults (significant channels were computed using uncorrected cluster based t-test  at each timepoints). In Insets: Comparison of topographies of CV and VQ for the three age groups in the duration where maximum number of channels are significant for VQ. Significant channels in these time-windows are marked in red. (D) time-lagged cross correlation between CV and VQ time-series, averaged across electrodes and subjects. The times of peak correlation are marked for each age group in both conditions. Inset picture shows the same for adults. Shaded areas mark S.E.M. across subjects for each cohort and conditions.

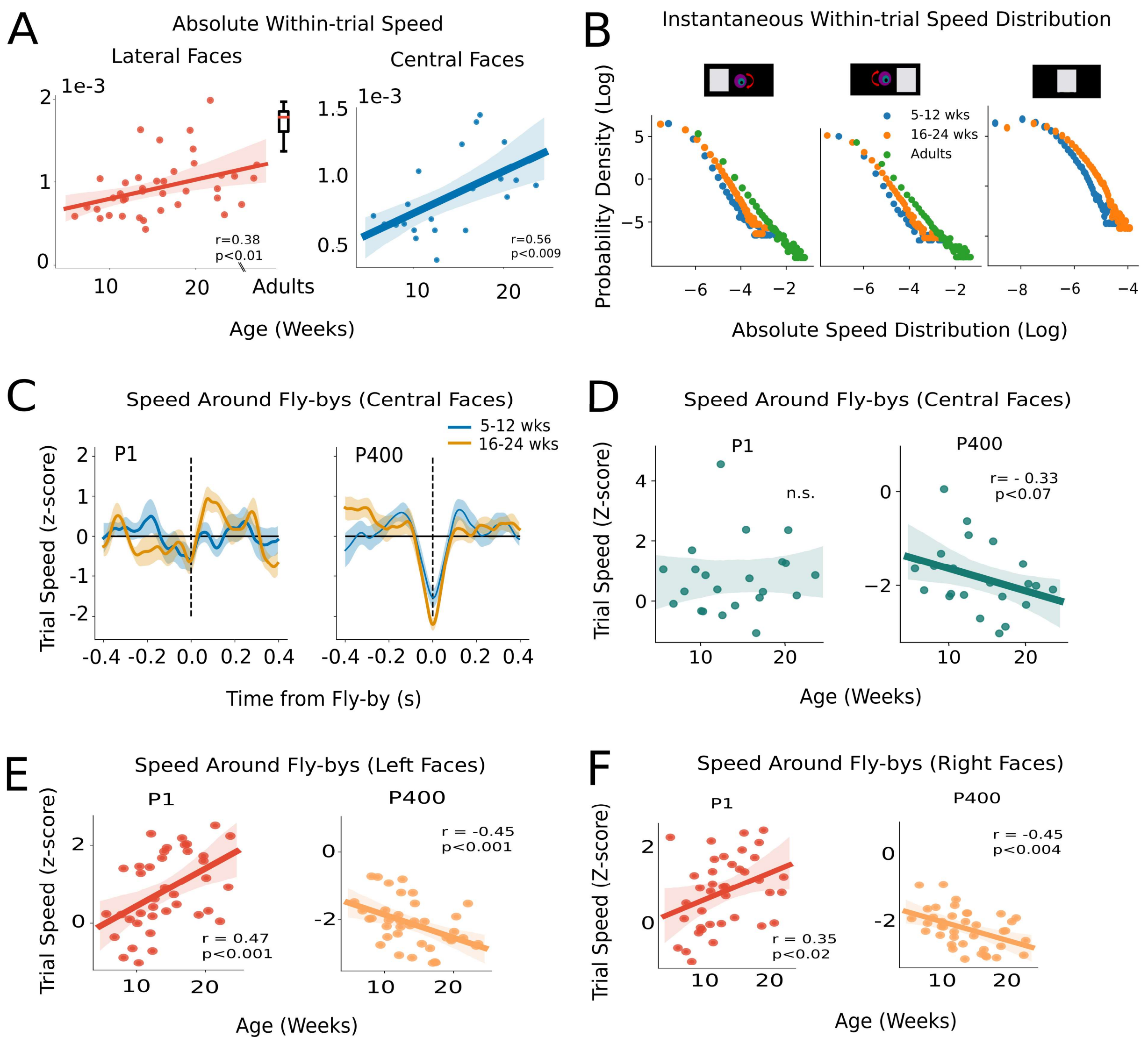


**Fig S7 Maturation of Within-trial Speeds. (A)** Absolute within-trial speed (averaged across all trials and times for each subject) for lateral and central faces (in left and right panels respectively). Absolute speed increases with age for each condition. Each dot represents one infant. Box plot represents the same metric for adults. **(B)** Instantaneous speed distributions (log-log plot) were heavy-tailed with adults and 16-24 week old infants being faster than 5-12 weeks infants for each condition. Each dot represents probability density for observing specific within-trial speed range at any time. **(C)**Relative within-trial speed profiles (z-scores) in the vicinity of P1 and P400-flybys for the faces presented on the central visual field. (**D-F**), Age trends on relative within trial speed in the vicinity of P1 (left panels) and P400 (right panels) for central **(D)**, left **(E)** and right **(F)** faces respectively. Each dot represents peak speed around the P1-flyby and lowest speed around P400-flyby in the 100-ms time-window surrounding the closest flyby. Solid lines represent least square fit to the data and shaded regions represent 95% confidence intervals. All r-values corrected for multiple comparisons with 1-tailed permutation tests.

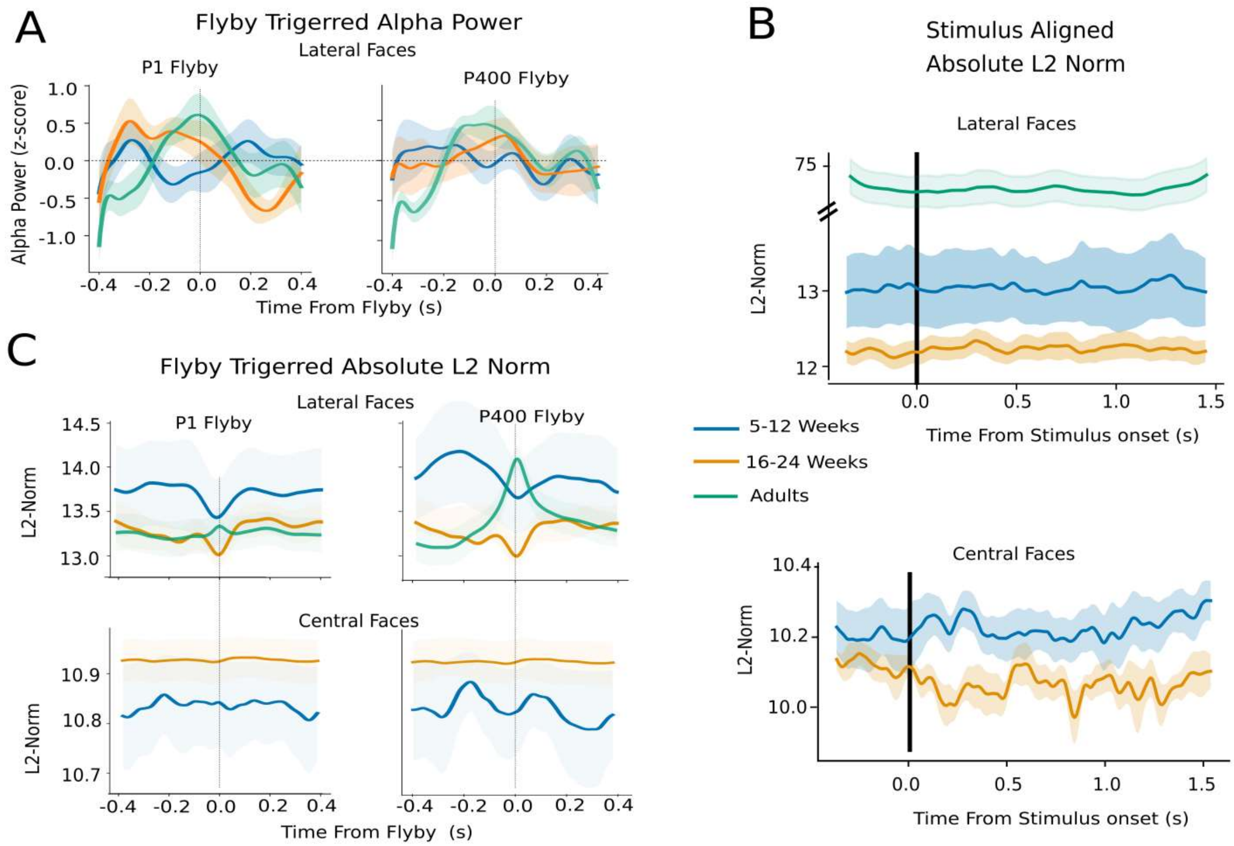


**Fig S8 Signal Strength or Signal to Noise Ratio. A)** Group averaged flyby triggered amplitude envelope of the alpha oscillatory component for young (5-12 weeks), old (16-24 week) infants and adults when they were presented with lateral faces. No significant modulation was observed for alpha power in the surroundings of P1 (left panel) or P400 (right panel) flybys. Power was averaged across all channels for each subject. **(B)** Broadband power or L2 Norm of the activation topographies (averaged across trials and across age-groups for lateral (top panel) and central faces (bottom panel). **(C)** Group averaged flyby triggered broadband power (or l2 norm) around the flyby to the ERP template P1 and P400( in left and right panels respectively) and for lateral and central faces (in top and bottom panels respectively). Shaded regions represents S.E.M. across subjects.
